# Mojave Desert microbial communities show high resistance and resilience over three years despite widespread plant mortality following the Dome Fire

**DOI:** 10.1101/2025.09.18.677208

**Authors:** Arik Joukhajian, M. Fabiola Pulido Barriga, Melanie J. Davis, Lynn C. Sweet, Sydney I. Glassman

## Abstract

1. High severity desert fires are uncommon but typically chart a new successional trajectory altering plant communities for at least 65 years. These aboveground vegetation shifts can have large implications for belowground microbial communities that maintain soil structure and nutrient cycling. High severity wildfires in forests or shrublands can severely reduce microbial species richness and biomass and alter microbiomes for decades but impacts on desert soil microbiomes are virtually unknown.
2. The 2020 Mojave Desert Dome Fire burned 43,273 acres of Eastern Joshua tree (Yucca jaegeriana) habitat, burning roughly 1 million trees. To track aboveground and belowground impacts of the Dome Fire, we established 9 plots (6 burned; 3 unburned) and sampled 4 subsamples per plot for 5 time points ranging from 2 weeks to 3 years post-fire. We measured initial ash depth as a proxy of soil burn severity and assessed plant mortality, plant richness, soil chemical characteristics, estimated soil microbial biomass with qPCR, and microbial richness and composition with Illumina MiSeq of 16S and ITS2 amplicons.
3. Belowground communities were highly diverse, containing 25,444 bacterial, 269 archaeal, and 6,683 fungal ASVs amplicon sequence variants (ASVs) or microbial taxa. We identified at least 65 plant species and saw 80% Eastern Joshua tree mortality in burned plots over three years, with reduced plant richness post-fire except an abundance of annual herbs at 1-year post-fire, yet the fire did not significantly reduce microbial biomass or richness at any time point.
4. Microbial communities for both bacteria and fungi showed small but significant changes, enriching for pyrophilous microbes in burned plots. We identified increases of pyrophilous microbes such as Tumebacillus, Massilia, Noviherbaspirillum bacteria and Pseudotricharina, Penicillium, Coniochaeta and Naganishia fungi.
5. Synthesis: We present the first comprehensive above and belowground examination following a natural desert wildfire including Archaea, Bacteria, and Fungi. Despite the widespread mortality of Eastern Joshua trees across 3 years, microbial biomass, richness, and community composition were mostly resistant to change, like microbial responses to low-intensity fast-moving grassland fires. Despite high resistance overall, wildfire still increased several pyrophilous bacterial and fungal taxa common after high severity shrubland and forest wildfires.

## Introduction

Drylands make up 46% of the terrestrial biosphere and are expected to expand to 56% by 2100 due to climate change (Cherlet et al., 2018). Increased mean temperatures, variability in precipitation, and invasive plants are increasing wildfire risks with long-lasting impacts on soil resources (Ravi et al., 2022). Frequently burned environments such as California chaparral shrublands (Hanes, 1971) and pine forests (Habeck & Mutch, 1973) have well understood vegetational responses to fire, but desert wildfires are infrequent and relatively understudied (Klinger et al., 2021; Schussman et al., 2006). Specifically, the Mojave Desert of Western North America has experienced increasingly large wildfires in recent decades (Brooks & Matchett, 2006; Klinger et al., 2021) with slow regenerating plants exhibiting century-scale secondary successional patterns (Abella, 2010). Moreover, fire impacts on desert soil microbiomes are nearly unknown (E. B. Allen et al., 2011), but could have critical implications for desert biogeochemical cycling (Homyak et al., 2016) and plant regeneration, since above- and below- ground communities are intimately linked (Van Der Heijden et al., 2008).

Desert plants like the iconic Joshua trees (Yucca brevifolia Engelm. and Yucca jaegeriana (McKelvey) L.W. Lenz) of the Mojave Desert are dependent on root-associated fungal communities for survival (M. F. Allen, 1989), yet these symbiotic associations are vulnerable to disturbances from fire (Dove & Hart, 2017). Arbuscular mycorrhizal fungi (AMF) are obligate symbionts associated with 70-80% of plant species (Brundrett & Tedersoo, 2018). AMF are critical partners for desert plants, especially Yucca (Titus, Titus, et al., 2002), since AMF increase plant access to water (Augé, 2001; Kakouridis et al., 2020) and enhance drought tolerance (Augé et al., 2015). Both the Western Joshua tree Yucca brevifolia (Harrower & Gilbert, 2021) and Eastern Joshua tree Y. jaegeriana (Joukhajian & Glassman, 2025) host diverse AMF from 8 Glomeromycotina families. Fungi can be either resistant (insensitive) or resilient (recover quickly to original status) to fire (Shade et al., 2012). Most long-term studies show that AMF have high post-fire resilience (Aguilar-Fernández et al., 2009; Bellgard et al., 1994; Espinosa et al., 2023) although fires can select for changes in AMF spore traits both immediately post-fire and 6-months later (Hopkins et al., 2024). Fires also damage biological soil crusts that house AMF and other root-associated fungi such as dark septate endophytes (Aanderud et al., 2018; Brianne et al., 2020), thus reducing sources of fungal spore dispersal (Warren et al., 2019). These indirect effects may lead to both short- and long-term restructurings of plant-associated fungal communities.

Although most fungal species are negatively impacted by fire, some pyrophilous “fire-loving” fungi that are rare or undetectable pre-fire massively increase in abundance post-fire (Fox et al., 2022). Several genera in Pyronemataceae are referred to as “phoenicoid fungi” for their abundant fruiting via ascocarps post-fire (Carpenter & Trappe, 1985). Studies in pine forests (Bruns et al., 2020; Caiafa et al., 2023; Glassman et al., 2016; Jacobson et al., 2004), shrublands (Pérez-Valera et al., 2017; Pulido-Chavez et al., 2023), boreal forests (Whitman et al., 2019), and eucalyptus forests (McMullan-Fisher et al., 2002; Warcup, 1990) highlight pyrophilous ascomycete species in the genera Pyronema, Neurospora, Penicillium and Aspergillus. In contrast, little is known about how pyrophilous fungi might respond in desert ecosystems, where high temperatures might pre-adapt common desert fungi like Naganishia (Lopez et al., 2024; Wei et al., 2022) and Alternaria (Malicka et al., 2022; Pombubpa et al., 2020) to survive wildfire. Studies on microbial community responses to desert fires are limited to controlled fires (Aanderud et al., 2019; X. Liu et al., 2000; Vega-Cofre et al., 2023; Zhang et al., 2025), where Dothideomycetes like Preussia lignicola often increased in abundance. However, one study after a natural desert wildfire in Mexico showed decreases in AMF richness post-fire (Chimal-Sánchez et al., 2015). As desert wildfires burn more acreage, understanding how symbiotic fungi are affected and whether pyrophilous fungi proliferate post-fire is essential for identifying successional trajectories and predicting ecosystem recovery.

Desert fires may also impact archaea and bacteria, which are overrepresented in desert chemical cycling pathways compared to other ecosystems (Makhalanyane et al., 2015). Abundant bacterial phyla in deserts include Actinobacteriota, Chloroflexota, Proteobacteria, and Planctomycetota. Archaea, particularly the phylum Crenarchaeota, are more abundant in deserts than other ecosystems (Ramond & Cowan, 2022). Wildfires in forested ecosystems broadly depress the phyla Acidobacteriota, Bacteroidota, and Verrucomicrobiota (Nelson et al., 2024; Rodríguez et al., 2018), but bacteria can also be pyrophilous. For example, members of Actinobacteriota like Conexibacter and Proteobacteria like Massilia and Noviherbaspirillum increased in abundance after a wildfire in chaparral shrublands (Pulido-Chavez et al., 2023) and forests (Caiafa et al., 2023; Soria et al., 2023). Firmicutes also grow in dominance post-fire (Enright et al., 2022; Pérez-Valera et al., 2017; Pulido-Chavez et al., 2023). Moreover, bacteria such as Arthrobacter and Massilia are known to persist in forests for up to 9 years after a wildfire and may be considered biomarkers of altered post-fire communities (Fernández-González et al., 2023). Rapid-responding prokaryotes are activated by rain pulses in the desert, allowing them to wait through periods of drought (Collins et al., 2014; Cowan et al., 2020). Yet, an increasingly variable precipitation regime may further hamper microbial communities, as decadal succession timelines in plants can disconnect nutrient cycling processes, leading to drainage of essential nutrients deep belowground (L. Wang et al., 2022). Thus, long-term sampling is critical to link plant and microbial responses to fires in the desert, where lag effects are common due to delayed plant mortality, slow nutrient turnover, and the rain-pulse nature of microbial activity (Maestre et al., 2021; Muñoz-Rojas et al., 2016; Vamstad & Rotenberry, 2010).

Fires can affect microbes directly via heat-induced mortality, or indirectly by changes in vegetation or soil chemical properties like increased pH (Certini et al., 2021; Hart et al., 2005). Fire intensity, or the heat transfer of fire, and fire severity, or the impact of fire on above- or belowground organisms, can shape wildfire impacts (Keeley, 2009). Fires affecting areas with lower or less dense aboveground biomass like annual grasslands and deserts are typically fast moving and low intensity (100-200°C) with low soil burn severity causing minimal impacts on associated soil microbes (E. B. Allen et al., 2011; Brooks, 2002; Glassman et al., 2023). In contrast, fires affecting areas with dense aboveground vegetation like shrublands and forests are typically slower moving, higher intensity (300-700°C), with higher soil burn severity with large reductions in soil microbial biomass and richness (DeBano et al., 1979; Dooley & Treseder, 2012; Nelson et al., 2022). Beyond heat-induced mortality, a permanent shift in aboveground plants may result in loss of important plant symbionts like AMF (Mummey & Rillig, 2006; Stinson et al., 2006). In August 2020, the Dome Fire burned 175 km^2^ (43,273 acres) of the Mojave Desert, burning roughly 1 million Y. jaegeriana trees (NPS., 2020). Although this fire was fueled by grasses and spread rapidly at low intensity, it was unusual in that it had high severity impacts on aboveground vegetation killing large tracts of long-lived perennial Y. jaegeriana trees and surrounding shrubland, causing concern over the long-term persistence of the species (Smith et al., 2023). This leads to a large knowledge gap, i.e. will microbial impacts be minimal like other fast moving low intensity fires, or large due to the indirect effects of widespread plant mortality?

Here, we investigate the short- and longer-term direct and indirect effects of the 2020 Dome Fire on above and belowground communities across five timepoints at 17 days, 1 and 8 months, and 1- and 3-years post-fire. We asked, 1) how did plant community composition and Y. jaegeriana survivorship change? 2) did archaeal, bacterial, and fungal richness and biomass decline? 3) were changes in bacterial and fungal community composition driven by direct or indirect effects? and 4) did pyrophilous microbes commonly seen in other fire-prone ecosystems proliferate? We hypothesized that the atypical severity of the fire on the long-lived perennial plant community would have a more deleterious impact on microbial communities despite being a fast-moving low-intensity fire, and that there would be lag effects in the response due to the delayed impacts of plant mortality and the rain-pulse driven nature of microbial responses in the desert. This would drive decreases in richness of bacteria, fungi, and AMF in particular, with minimal impact on the highly resistant archaea (Makhalanyane et al., 2015), but might lead to increases in pyrophilous microbes like fungal Dothideomycetes, which were common after prescribed burns in deserts (Zhang et al., 2025), and bacterial phyla Firmicutes and Proteobacteria, which are common after fire in many ecosystems (Enright et al., 2022; Pérez-Valera et al., 2017; Pulido-Chavez et al., 2023).

## Materials and Methods

Site description: We sampled in Mojave National Preserve, California, USA, around the Cima Dome (35.264777°, -115.516038°), home to the densest Y. jaegeriana stand, where the Dome Fire burned 175km from August 15-24 2020 (Fig. 1a). Our study area had a mean elevation of 1361m, in a region with bimodal summer and winter rains for at least the last century (Tagestad et al., 2016). Vegetation in the nearby unburned area consisted of Y. jaegeriana shrublands, including Y. baccata, Y. schidigera and annual and perennial grasses like Bouteloua spp., Bromus spp., and Hilaria jamesii. Shrubs included Ambrosia salsola, Coleogyne ramosissima, Ephedra nevadensis, and Lycium spp.

**Figure 1.**
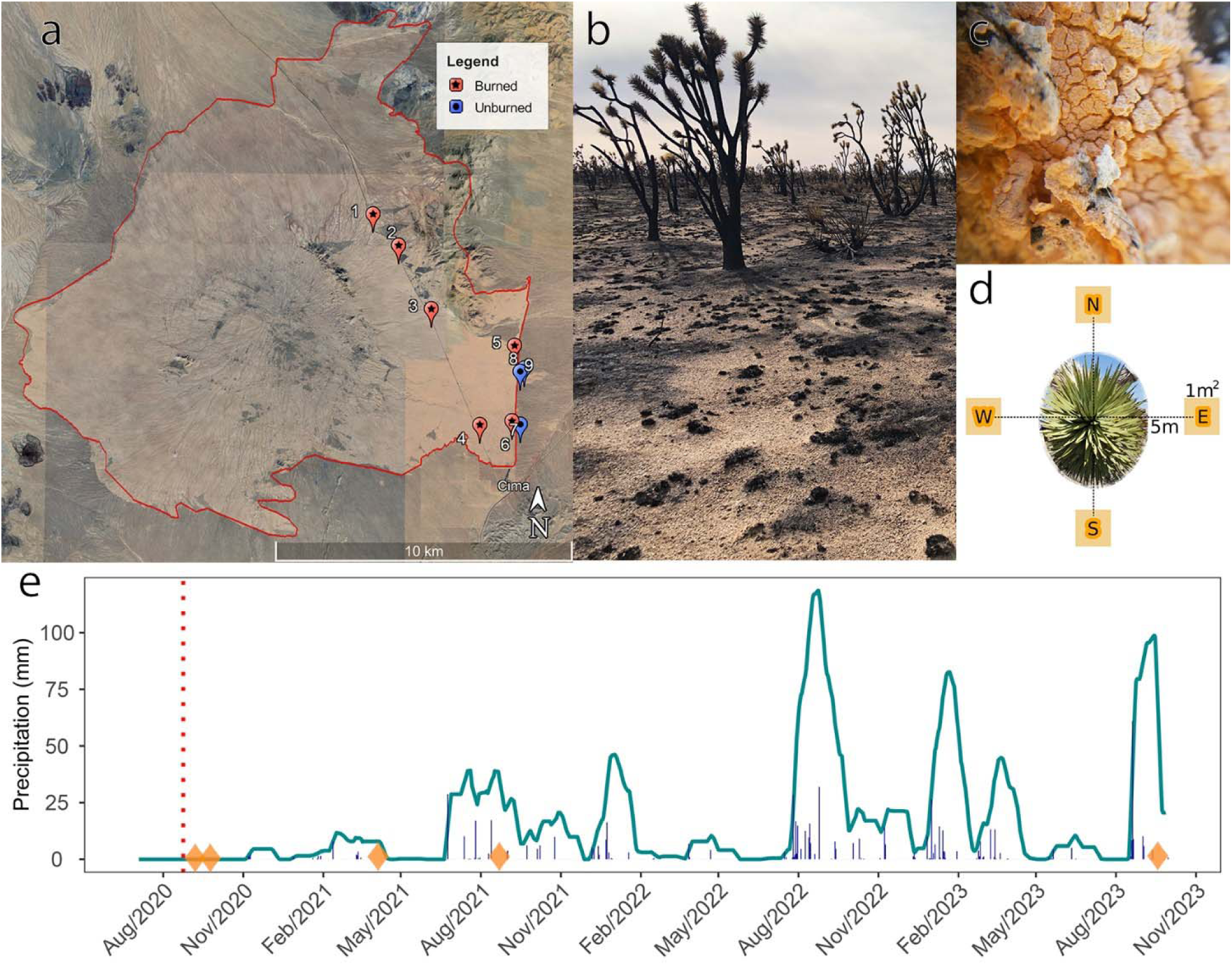
a) A map of our 9 plots, with the Dome Fire perimeter in red. Unburned plots (7-9) are east of the boundary on Morning Star Mine Rd. b) Y. jaegeriana and burnt soil in the Dome Fire burn scar 17 days after the end of the fire. c) Neurospora discreta growth on Y. jaegeriana bark at 8-months post-fire. d) A schematic demonstrating the 4 1m^2^ subplots within the 9 plots. e) Precipitation at the Mojave Mid Hills station. Bars indicate recorded precipitation per month and the blue line indicates a rolling sum of the prior 4-months of precipitation. The red dash marks the Dome Fire, while orange diamonds indicate our 5 sampling dates from 17-days, 1-month, 8-months, 1-year and 3-years post-fire.

Experimental design and soil collection: In September 2020, 17-days after the fire was extinguished, we established nine plots (6 burned and 3 unburned; Fig. 1a), each centered around a singular live or dead Y. jaegeriana tree (Fig. 1b). Many trees were highly colonized by the fungus Neurospora discreta underneath exposed bark (Fig. 1c), which were identified by culturing and sequencing the genome. Within each plot, we established four 1m^2^ subplots 5m from the focal tree in each of the cardinal directions for a total of 36 subplots (Fig. 1d). During the first sampling timepoint, we estimated soil burn severity by averaging three ash depth measurements from within each 1m^2^ subplot and estimating percent char at each subplot. We collected soil from the top 10 cm beneath the ash or organic layer with a releasable bulb planter cleaned between subplots with 70% ethanol to reduce contamination and homogenized 2 soil cores from each sub-plot. We collected soils at five time-points: September 10 and 24 2020, April 15 and September 9, 2021, and September 19, 2023, corresponding to 17-days, 1-month, 8-months, and 1 and 3-years post-fire. Our 3-year sampling occurred one month after Tropical

Storm Hilary brought the largest 1-day precipitation event in our study period, registering at 61mm of rain on August 20, 2023 (Fig. 1E). We collected soils from all 9 plots at all time points except for the second timepoint at 1-month post-fire where we were only able to collect soils from 7/9 plots including 1 unburned plot (plot #8). Soils were collected and stored on ice and transported to the University of California, Riverside (UCR), where we homogenized and sieved (2mm) within 24hrs of collection. We immediately analyzed soils for gravimetric soil moisture content and stored a portion at -80°C for molecular analysis for all time points. For 3 time points (17 days, 8 months and 3 years post-fire), we performed the KCl extraction method on fresh soil (Carter & Gregorich, 2007) and assessed ammonium, phosphate, and NO!il⁻ at the UCR Environmental Science Research Lab. We measured pH on air-dried soil by making a slurry of 10g of soil and 20mL of distilled water and measuring with a VisionPlus pH6175 meter (Jenco Instruments Inc., San Diego, CA) after one minute of shaking to mix.

Plant communities: We surveyed plant species and Y. jaegeriana mortality at 17-days, 1-year, and 3-years post-fire at each plot. We used a point centered quarter survey method (PCQM), which is designed to capture density in a random sample of plants (Cottam et al., 1953). For each plot, we located the nearest Y. jaegeriana to the center point of the sampling area from hereon referred to as the sentinel tree. At this sentinel tree we performed a Rapid Assessment (CNPS-CDFW) survey of the surrounding 20-meter radius, which included perennial plant richness and ocular estimates of cover, center point coordinates in UTMs center point coordinates in UTMs (Zone 11, WGS 84) and cardinal photos. From the sentinel, we identified each cardinal direction and found the nearest tree in each quadrant. Each site had a minimum of five trees (sentinel, NE, NW, SE, & SW). During the first sampling, we noted pre-fire mortality of trees by the sole presence of ash on the ground (biomass consumed), fire-related mortality via presence of char on the trees and absence of green leaves, and at subsequent samplings, noted resprouting or persisting green leaves as signs of survival. Then at each tree we took note of living and dead terminal ends of the tree, height of the tallest leaf blade, herbivory, and char.

DNA Extraction and Sequencing: We extracted DNA from 0.25g of frozen soil with Qiagen DNeasy Power Soil Pro Kits, modified by replacing 100 μL of the initial C1 solution with 100 μL ATL and incubating overnight at 4°C to improve yields. We PCR amplified DNA with the primer pair 515F-806R targeting the V4 region of the 16S rRNA gene of bacteria and archaea (Caporaso et al., 2011) and the 5.8S-ITS4 primer pair to amplify the ITS2 region (Taylor et al., 2016) of the Internal Transcribed Spacer, the internationally recognized barcode for fungi (Schoch et al., 2012). We attached dual-index primers (DIP) with barcodes and adaptors onto amplicons with a second PCR step (Kozich et al., 2013). To amplify ITS2, we mixed 5µL DNA with 0.5 μL of the 5.8S and ITS4 primers (10 μM), to amplify 16S, we mixed 1 μL DNA with 0.5 μL of the 515F and 806R primers (10 μM), then added 12.5μL AccuStart II PCR Tough Mix (Quantabio, Beverly MA, USA), and added Ultra-Pure Sterile Molecular Biology Grade water (Genesee Scientific, San Diego, CA, USA) up to a 25 µL reaction. We cleaned PCR1 amplicons with AMPure XP magnetic beads (Beckman Coulter Inc.) following manufacturer instructions then attached DIP-barcodes in PCR2 with the following mix: 2.5 μL of 10 mM DIP PCR2 primers, 6.5 μL of ultrapure water, 12.5 μL of Accustart II PCR ToughMix and 1 μL PCR1 product. See Table S1 for PCR thermocycler settings.

We pooled 16S and ITS2 PCR products separately based on band-strength then cleaned the pools with Ampure XP magnetic beads, included negative controls for DNA extractions and PCRs, and included the Microbial Standard II (Zymobiotics, Orange, CA) mock community in 16S and ITS2 pools. We then assessed amplicon pool size and intensity with the Bioanalyzer before pooling at a 2:3 ratio of 16S to ITS2 to account for the short amplicon preference of Illumina sequencing and the typically longer ITS2 amplicons. We sequenced with Illumina MiSeq 2x300bp at the UCR Institute for Integrative Genome Biology across two sequence libraries.

Biomass: We estimated sequence copy number as a proxy of microbial biomass with quantitative polymerase chain reaction (qPCR) for either the 16S for archaea and bacteria using the Eub338/Eub518 primer pair (Fierer et al., 2005) or the 18S FungiQuant primers for fungi (C. M. Liu et al., 2012). Each 10 μL qPCR reaction contained 1 μL DNA, 5 μL 1X iTaq SYBR mix (Bio-Rad, Hercules, CA), 0.4 μL (10 μM) of each primer, and 3.2 μL ultra-pure water with thermocycler settings in Table S1. We ran samples in triplicate on the CFX Opus 384 Real-Time PCR System (Bio-Rad, Hercules, CA). We used standards with a known concentration in a dilution series to determine sample starting concentration and calculated gene copy numbers per gram of soil with the following equation: Copy number = (Starting concentration (ng/ μL) * (1 μL in reaction) * 6.022 * 10^23 (Avogadro’s constant)) / (Length of standard (bp) * 660 ng (average weight of a base pair)* 10^9 (average number of cells per g of soil).

Bioinformatics and statistical analysis: We analyzed Illumina Miseq data using QIIME2 (Bolyen et al., 2019) to produce tables of Amplicon Sequence Variants (ASVs). We removed forward and reverse adaptors using the QIIME2 cutadapt trim-paired plugin (Martin, 2011), demultiplexed, and denoised using DADA2 (Callahan et al., 2016) with the denoise-paired plugin based on the quality of forward reads to ensure median quality scores over 30. We assigned taxonomy of ITS2 sequences using the dynamic database in UNITE v.9 (Abarenkov et al., 2010), and 16S sequences using the database SILVA v.138 (Quast et al., 2013). We then exported the ASV table to R ver. 4.1 (R Core Team, 2020), where we normalized the sequencing depth per Kingdom (638 for archaea, 4,256 for bacteria, and 6,874 for fungi) to account for uneven sequencing depth (Schloss, 2024), and estimated alpha diversity metrics with the BiodiversityR package (Kindt, 2024). We used qiime2R (Bisanz, 2018) to analyze QIIME2 outputs in R and phyloseq (McMurdie & Holmes, 2013) to generate relative abundance barplots and line graphs from composition data, and “Probable” or “Highly Probable” guild data of fungi from FUNGuild (Nguyen et al., 2016) using FungalTraits (Põlme et al., 2020). We used generalized linear mixed (GLMM) effects models to test the impact of fire and time on microbial richness and biomass with plot as a random effect using lme4 (Bates et al., 2015). We checked the best fit distribution of each variable with the “fitdistr” function in the MASS package (Ripley et al., 2024).

To identify shifts in plant communities, we used estimates of plant cover abundance to compare overall relative abundance of plant families between the burned and unburned plots at 17-days, 1-year, and 3-years post-fire. At the same timepoints, we summed total counts of living and dead Y. jaegeriana across burned and unburned plots, which included previously felled trees in unburned plots, and plotted the percentage of total surviving trees per treatment. At each individual timepoint where plants were measured, we used ANOVA to test fire effect on plant richness and used the mantel function within the vegan package (Oksanen et al., 2008) to correlate plant and microbial composition.

To identify drivers of microbial community compositional changes, we used the vegan functions “adonis2” and “betadisper” to identify when changes were due to increased heterogeneity across a group. To compare the impacts of soil properties or plant richness on microbial communities, we visualized beta diversity using non-metric multi-dimensional scaling (NMDS) plots of Bray-Curtis dissimilarity with the vegan function “metaDMS”, overlaid continuous variables with the “envfit” function, and adjusted p-values with the Bonferroni method using “p.adjust” for repeated measurements across time for the time points with matching microbial composition, soil chemical, and plant richness data. After determining that days post-fire was not a significant driver of community dissimilarity for each subset of samples and timepoints, we combined samples across time. We separated the effect of spatial autocorrelation with the vegan function “varpart” using a distance matrix, a PCA of environmental variables, and each Bray-Curtis dissimilarity matrix. To identify pyrophilous microbes, we used the Deseq2 package (Love et al., 2014) to identify genera with significant log2fold changes (alpha of 0.01) in burned relative to unburned plots at each time point (excluding the 1-month timepoint due to the low number of unburned samples).

## Results

Fire impact on plant mortality, richness, and diversity: At 17-days post-fire, burned subplots had a mean ± standard error ash depth of 0.37 ± 0.07 cm (Fig. 2A) and 43.33% ± 4.37 char percentage (Fig. 2B) whereas unburned plots had none. In unburned plots, Y. jaegeriana survivorship was stable at roughly 83% across all time points (Fig. 2C). In contrast, in burned plots, Y. jaegeriana survivorship decreased across time with percentage of trees having some green leaves reducing from 86% at 17-days to 50% at 1-year to only 20% at 3 years post-fire (Fig. 2C). Fire significantly reduced plant richness from 12.67 ± 0.33 in unburned to 2.32 ± 0.94 in burned plots at 17-days post-fire (Table S2; Fig. 2D). However, emergence of annual plants led to burned plots (8.6 ± 2.7) having similar richness to unburned plots (12 ± 1.5) at 1-year post-fire (Fig. S1). Yet this change was temporary as plant richness was significantly lower in burned (6 ± 0.6) than unburned plots (11.6 ± 2.3) at 3 years post-fire. Fire initially obliterated plant biomass reducing it from 50% ± 2.78 cover in unburned to nearly undetectable at 0.23% ± 0.09 in burned plots at 17 days post-fire (Table S2; Fig. 2E). Across three years, we observed 65 plant species from 47 unique genera across 19 plant families in burned and unburned plots with large changes in composition at 1- and 3-years post-fire with a notable loss of Cactaceae in burned plots (Fig. 2F). Several plants were first detected in burned plots 1-year post-fire, mainly comprised of native annual and perennial grasses and herbs with only one non-native plant Erodium cicutarium, which had minimal cover (Table S3).

**Figure 2.**
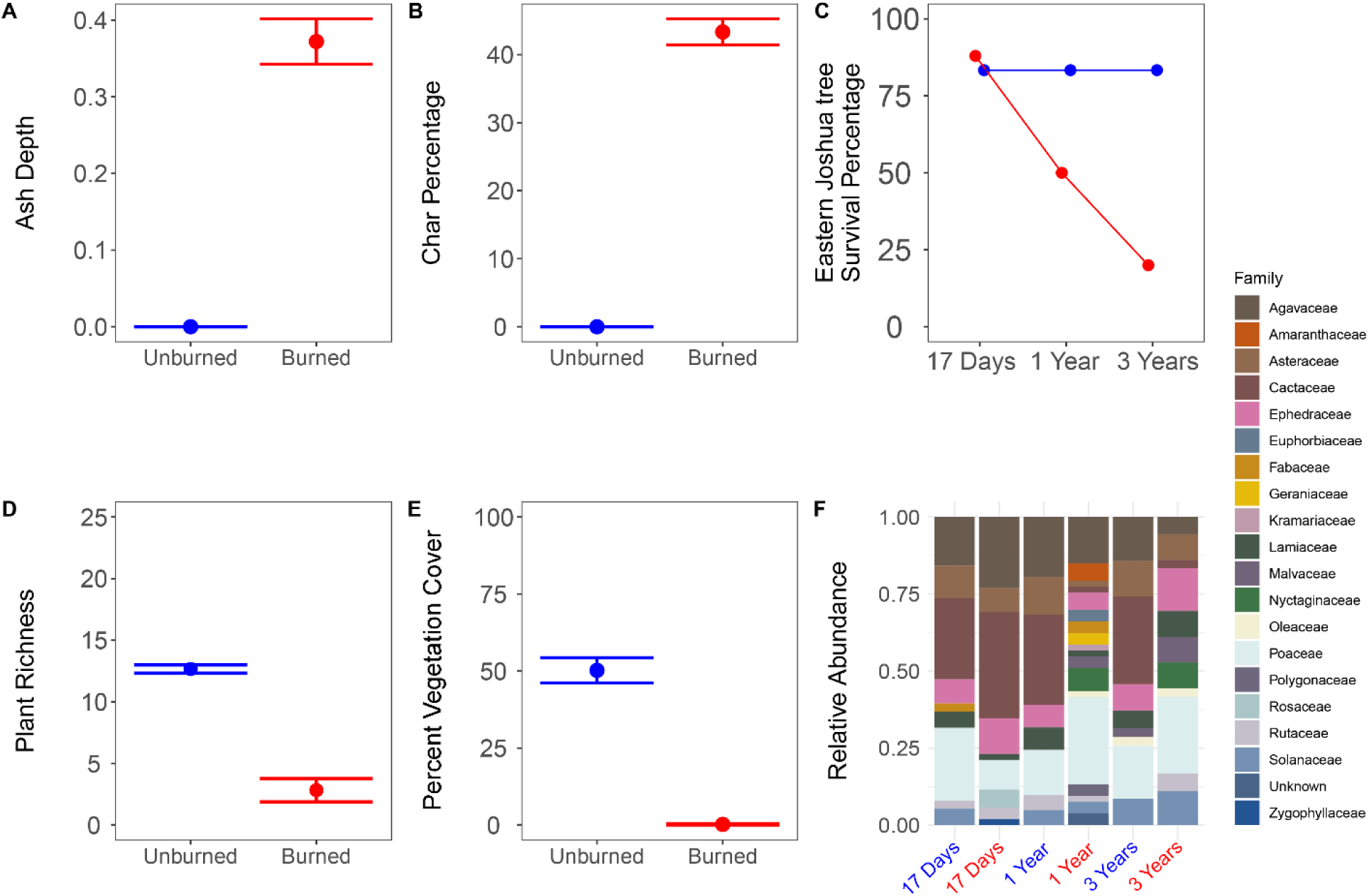
Mean and standard error bars of: A) Soil burn severity via ash depth in cm. B) Char percentage on the soil surface measured for each subplot at 17-days post-fire. C) Percentage of Y. jaegeriana assessed as “living” in 6 burned plots and 3 unburned plots, including previously felled trees. Mean and standard error bars of: D) Plant species richness E) Percent cover of vegetation measured for burned versus unburned plots at 17-days post-fire. F) Plant family relative abundance across unburned (blue) and burned (red) plant families assessed post-fire at 17 days in September 2020, at 1 year in September 2021, and 3 years in September 2023.

Fire impact on soil: There were significant effects of fire, time, and fire by time interactions on pH such that fire significantly increased soil pH in burned plots overall and at each time point individually except 8 months (Table S4, Fig. S2A). Fire did not significantly affect phosphate but time did, such that overall mean phosphate decreased from 17-days to 8-months and increased from 8-months to 3-years post-fire across burned and unburned plots (Table S4, Fig. S2B). Fire did not have a significant direct effect on ammonium but time did such that ammonium decreased across time in burned and unburned plots (Table S4, Fig. S2C). There was also a significant fire by time interaction such that ammonium was greater in burned than unburned plots at 8-months and 3-years post-fire (Table S4). Both time and its interaction with fire had significant effects on nitrate and nitrite such that they increased over time but nitrate and nitrite were higher in burned than unburned plots at 3-years post-fire (Table S4, Fig. 2D).

Fire impacts on microbial richness and biomass: We generated 18,338,870 ITS2 and 14,963,273 16S reads across two libraries. After quality filtering and assigning taxonomy, we had 269 archaeal, 25,444 bacterial, and 6,683 fungal ASVs across our 172 samples from 5 timepoints across 3 years. Fire did not have the direct effect of reduction in ASV richness of bacteria, fungi, archaea or AMF (Table S5, Fig. 3). Furthermore, soil burn severity also did not significantly correlate with richness of fungi (Table S5, Fig. S3) or bacteria (Table S5, Fig. S4) any timepoint.

**Figure 3.**
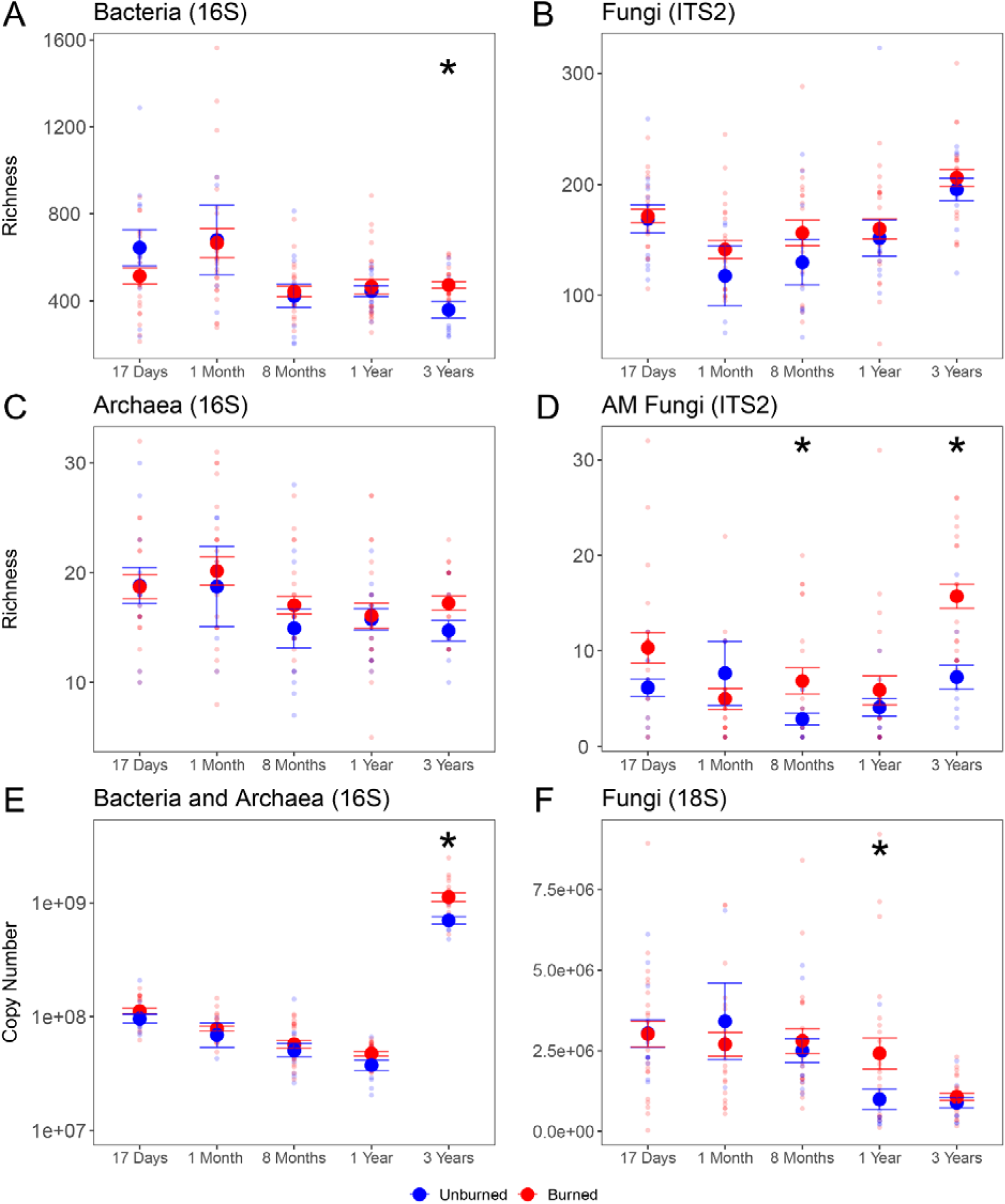
Mean ± standard error of richness and copy number in burned and unburned plots (red and blue, respectively) at each timepoint. Richness of bacteria, fungi, archaea, and arbuscular mycorrhizal fungi are represented by A,B,C, and D respectively. Bacterial and archaeal 16S and fungal 18S are represented by E and F, respectively. Asterisks indicate significant differences between means of burned and unburned plots at a given timepoint. Unburned plots at 1-month only have 4 total samples, causing high variance.

However, there were significant time by fire interactions such that burned plots showed greater bacterial richness at 3-years (Table S5, Fig. 3A), and AMF richness at 8-months and 3-years (Table S5, Fig. 3D) relative to unburned plots (Table S5). Fire also did not have a significant direct effect on 16S or 18S copy number (Table S6). However, there was a significant fire by time interaction such that burned relative to unburned plots had higher copy numbers of 16S at 3 years (Table S6, Fig. 3E) and 18S at 1 year (Table S6, Fig. 3F). Finally, there was a significant effect of time on both 16S and 18S such that across burned and unburned plots, bacterial biomass decreased from 1 month to 1-year post-fire but increased at 3 years and fungal biomass decreased at 3 years (Table S6). When using model selection to determine if precipitation or soil moisture were better predictors than time alone, we found that time was the best predicter of bacterial richness and biomass (Table S7) and fungal richness (Table S8). However, precipitation in the last 90 days was a better predictor of fungal biomass when compared to time and soil moisture (Table S8).

Fire and time impacts on community composition: Fire had small but significant impacts on fungal and bacterial community composition at all 5 time points and on archaeal composition at 8 months and 3 years (Table S9). Community heterogeneity (beta-dispersion) temporarily increased for archaea, bacteria, and fungi for up to 1-month, but homogenized afterwards, such that burned plots became more alike while remaining distinct from unburned plots (Table S9).

Drivers of microbial community composition over time and links to plants and soil properties: Char percentage was a significant driver of bacterial (Fig. 4A), archaeal (Fig. 4B), and fungal composition (Fig. 4C). Bacterial community dissimilarity was driven primarily by pH, followed by soil moisture, char percentage, nitrate and nitrite, and ammonium (Fig. 4D). Archaeal community dissimilarity was driven primarily by soil moisture, followed by pH, char percentage, and nitrate and nitrite. Fungal community dissimilarity was driven primarily by char percentage, followed by pH, and plant richness. While plant richness was a significant driver of fungal community composition, plant community composition did not correlate to shifts in archaeal (r = 0.02, p = 0.42), bacterial (r = 0.03, p = 0.48), or fungal (r = 0.03, p = 0.33) communities. Although plots across our two treatments were spatially clustered, we note that environmental signals do not appear to be artefacts of spatial autocorrelation for any timepoints except for 17-days in bacteria and archaea but not fungi (Table S10).

**Figure 4.**
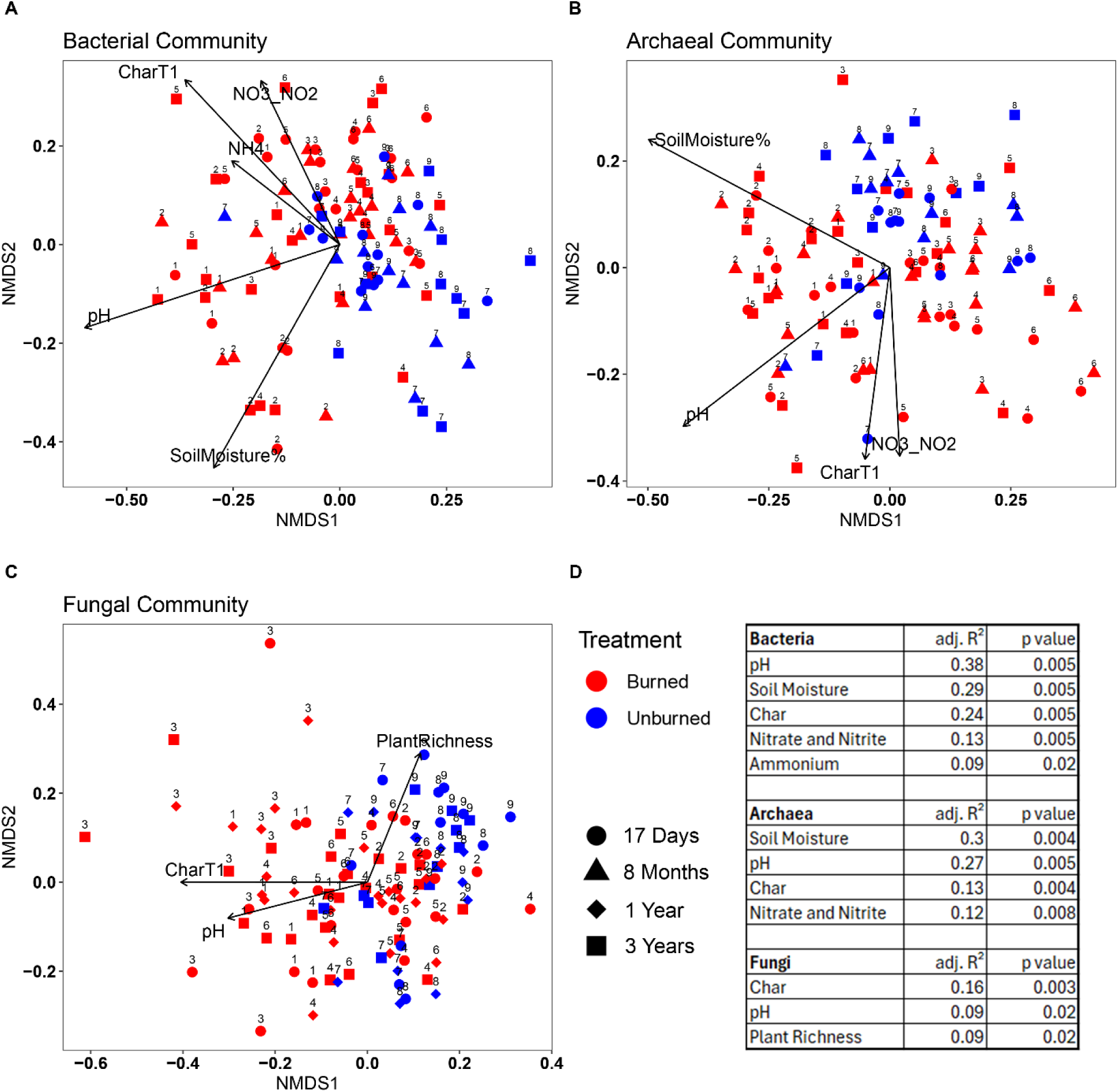
Bray-Curtis dissimilarity plotted by non-metric multidimensional scaling with environmental variables overlaid. CharT1 is the percent char of each subplot as measured at 17-days post-fire. NH4 is Ammonium, and NO3_NO2 is Nitrate and Nitrite. Arrow-length corresponds to the strength of each correlation, independent of the other factors. Bacterial (A) and archaeal (B) communities include samples from 17-days, 8-months, and 3-years post-fire. Fungal (C) communities include samples from 17-days, 1-year, and 3-years post-fire. Each point is labeled with the Plot number it was sampled from (Plot 1-6 are burned, Plot 7-9 are unburned). D) Significant variables (p < 0.05) and their adjusted R^2^.

Impact of fire on specific prokaryotic taxa and emergence of pyrophilous bacteria and archaea: Bacterial communities were dominated by the phyla Actinobacteriota, followed by Proteobacteria and Acidobacteriota (Fig. S5A). Actinobacteriota increased in burned plots at 1 month and 3 years, while Proteobacteria was consistently higher in burned plots, with a near 7% shift at 1-year (Fig. S5A). Bacteroidota and Firmicutes were more abundant in unburned plots, whereas Acidobacteriota were more abundant in burned plots. Archaea (6-9% of 16S reads) were more abundant in burned plots (Fig. S5A) primarily represented by the Crenarchaeota genera Candidatus Nitrocosmicus and Ca. Nitrososphaera (Fig. 5B). Cyanobacteria were more abundant in unburned plots during the first month post-fire with Tychonema dominating abundance (Fig. S5B). The most abundant genera in burned and unburned plots alike were Rubrobacter (Actinobacteriota), RB41 (Acidobacteriota), and Ca. Nitrososphaera (Crenarchaeota) (Fig. S5B).

**Figure 5.**
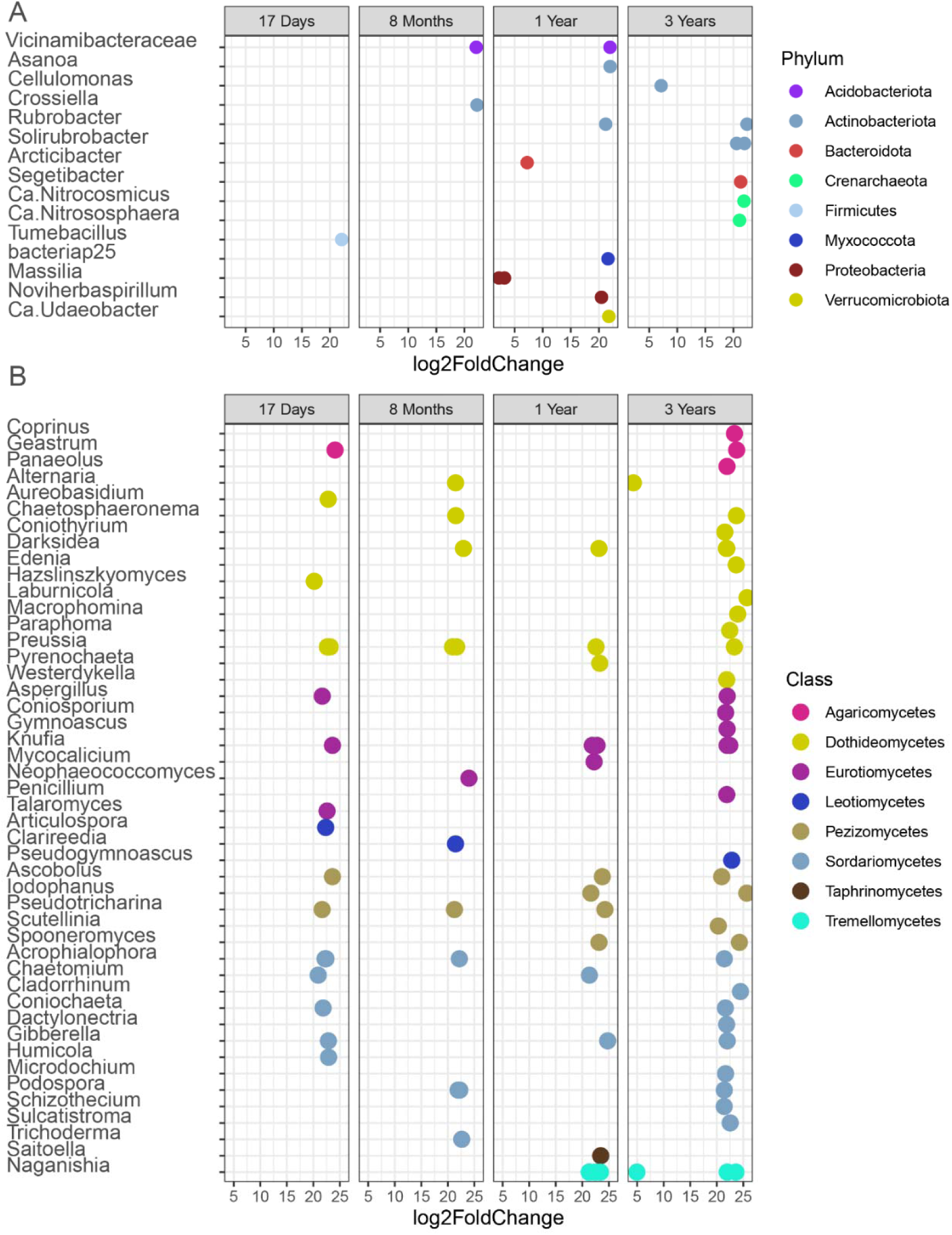
Microbial ASVs with increased differential abundance in burned plots at 4 timepoints, sorted by A) phylum for prokaryotes or B) class for fungi. Greater log2fold Change on the x-axis indicates more abundance in burned plots relative to unburned plots. A full list of archaeal, bacterial, and fungal ASVs, including ASVs responding negatively to fire, can be found in the supplemental materials (Table S11-14) as overlapping points may obscure multiple ASVs within one genus.

There were 2 archaeal and 17 bacterial genera that responded positively to fire including established pyrophilous microbes like Massilia and Noviherbaspirillum, and overall 1 ASV at 17-days, 2 at 8-months, 16 at 1-year, and 8 at 3-years post-fire (Fig. 5A; Table S11). The phyla with the most ASVs showing increased differential log2fold abundance in burned relative to unburned plots were Actinobacteriota with 8 ASVs, Proteobacteriota with 7 ASVs, and Firmicutes with 3 ASVs despite the overall lower relative abundance of the phylum Firmicutes in burned plots. Most bacterial ASVs showing positive differential log2fold abundance in burned plots were undetectable in unburned plots at that same timepoint (Fig. 5A). In contrast, only 7 ASVs negatively responded to fire across 3 years (Table S12).

Impact of fire on specific fungal taxa and emergence of pyrophilous fungi: Fungi were dominated by phylum Ascomycota, ranging from 81-88% of reads in burned plots and 84-94% of reads in unburned plots (Fig. S6A). Basidiomycota were slightly more abundant (9-16%) in burned than in unburned plots (5-14%; Fig. S6A). Basidiomycota were dominated by the mushroom-forming genera Lepiota, Montagnea, Disciceda, and Geastrum, and the basidiomycete yeast Naganishia (Fig. S6B). The overall most abundant fungal genus in both burned and unburned plots was Alternaria (Fig. S6B). Glomeromycotina (AMF) composed < 1% of the reads, except for 3-years post-fire where they in relative abundance in burned plots (Fig. S7).

There were 88 fungal genera that responded positively to fire including established pyrophilous taxa Naganishia, Coniochaeta, and Aspergillus which combined had a total of 26 ASVs at 17-days, 20 ASVs at 1-month, 21 ASVs at 8-months, 34 ASVs at 1-year, and 62 ASVs at 3-years (Fig. 5B; Table S13). The fungal classes with the most ASVs showing positive log2fold differential abundance in burned relative to unburned plots were Sordariomycetes (50 ASVs), Dothideomycetes (48 ASVs), and Eurotiomycetes (24 ASVs) within Ascomycota and Tremellomycetes (10 ASVs) and Agaricomycetes (6 ASVs) within Basidiomycota. In contrast, only 35 ASVs negatively responded to fire (Table S14).

## Discussion

Here, we present the first comprehensive examination of direct and indirect effects of a large desert wildfire on above and belowground communities including the iconic Y. jaegeriana, surrounding desert plants, Archaea, Bacteria, and Fungi. Despite ∼80% Y. jaegeriana mortality, archaeal, bacterial and fungal biomass and richness were highly resistant to wildfire at all time points ranging from 2 weeks to 3 years post-fire. Interestingly, by 3 years post-fire, AMF and bacterial richness and 16S biomass were elevated in burned relative to unburned plots. There were minor but significant impacts of fire on microbiome composition that persisted across the 3 years across the 3 domains, leading to the emergence of several pyrophilous microbes typically seen after high severity wildfires in shrublands and forests, such as the bacteria Massilia and Noviherbaspirillum, the archaea Ca. Nitrososphaera and Ca. Nitrocosmicus, and the fungi Penicillium, Coniochaeta, Aureobasidium, Alternaria, and Naganishia. Microbial communities were explained by various environmental factors including char percentage, but only fungal composition could be linked to plant richness. Together, these findings reveal initial resistance to fire and long-term resilience in microbial richness and biomass but small fire-induced shifts in community composition reflecting domain-specific environmental drivers and links between plant and fungal communities.

Dome Fire impacts on plants: The Dome Fire led to persistent and widespread mortality of Y. jaegeriana, which did not recover, but rather became surrounded by herbs, forbs, and grasses. In the Mojave, other post-fire plant communities have shown little recovery to unburned composition after decades (Abella, 2009) and in some cases, turnover into new and persistent community types (Abella et al., 2021). Yucca jaegeriana mortality was severe, increasing from 14% to 80% over the three-year span, similar to findings for Y. brevifolia (DeFalco et al., 2010). The delayed mortality of this species emphasizes the long-term vulnerability of Mojave plant communities to additional stressors such as post-fire drought (DeFalco et al., 2010). Annual plant community dynamics in deserts are influenced by the amount and timing of precipitation, including the abundance of exotic annual grasses that fluctuates year-to-year (Brooks & Berry, 2006), and post-fire plant communities are no exception. At 1-year post-fire, the rise in plant richness was driven by an abundance of annual grasses. While small patches of exotic grasses such as Bromus madritensis at 17-days post-fire and Erodium cicutarium at 1-year post-fire were present, native annual herbs and grasses led to increased plant richness in burned plots at 1-year. However, most annual herbs were ephemeral in their site occupancy, not persisting to 3-years post-fire, leading to the reestablishment of native perennial herbs, grasses, and shrubs and reduced plant richness in burned compared to unburned plots at 3 years post-fire. While deserts are vulnerable to exotic grasses increasing wildfire risks, Bromus madritensis was the only exotic annual grass present in unburned plots in 2020. Unburned plots saw an increase in perennial rather than annual grasses, and these similarly reduced in cover by 3-years. While shrubs are essential resource islands for water and nutrients in deserts (Titus, Nowak, et al., 2002), it remains unclear whether the reemergence of shrubs as the dominant cover in burned plots will drive convergence between burned and unburned plant communities.

Dome Fire impacts on plant associated fungi and AMF: While there were no direct effects of fire on AMF richness, changes in plant richness and composition were correlated with both AMF richness and overall fungal composition. In contrast to a different desert wildfire in Mexico, where AMF richness declined after fire (Chimal-Sanchez 2015), here, AMF richness increased over time in burned relative to unburned plots, with increased pH as a result of the burn potentially driving AMF community changes as it does globally (Davison et al., 2021). The abundance of AMF spores are heavily reduced in interspaces between Mojave plants (Titus,

Nowak, et al., 2002) so dominance of dark septate endophytes like Alternaria in these regions is expected, especially with the region’s history of cheatgrass invasion which increases colonization of dark septate endophytes (Gehring et al., 2016). Plant richness had a strong effect on overall fungal community composition, as one burned plot (#3), consistently ranked higher in plant species richness (Fig. S1) and showed much greater fungal community dissimilarity to other burned samples. This clustering effect was not present in bacterial or archaeal communities (Fig. 4).

Dome Fire impacts on soil chemical properties: Fire led to increased soil pH and inorganic nitrogen, aligning with established post-fire trends (Agbeshie et al., 2022). Ash deposition drove soil pH to become more basic in burned plots at nearly all timepoints, despite an ash depth of only <1cm compared to much higher soil burn severity of chaparral shrublands consisting of ∼12 cm ash (Pulido-Chavez et al., 2023). Similar to the chaparral shrublands that burned at higher soil burn severity (Pulido-Chavez, 2023), ammonium was elevated in burned compared to unburned plots at both 8 months and 3 years post-fire, whereas nitrate and nitrite levels were comparable in burned and unburned plots until year 3, when they became elevated in burned compared to unburned plots. Ammonium, which volatilizes in much greater amounts compared to nitrate and nitrite (Benedict et al., 2017), appears to have evenly deposited on burned and unburned plots, yet was depleted in unburned plots over the years, potentially through AMF more readily transferring it to plants (Meng et al., 2015). Meanwhile, heavy rainfall spurring microbial activity may have driven oxidization of this excess ammonium to nitrate in burned plots at 3-years post-fire (Z. Wang et al., 2020), resulting in several high readings of nitrate and nitrite in burned plots (Fig. S2D). While phosphate only ephemerally increases post-fire (Neary et al., 1999), it exhibited an ephemeral drop at 8 months and a rise at 3-years post-fire due to several outlier samples, potentially in line with nitrate and nitrite similarly showing “hotspots” of activity (Barnes et al., 2024).

Dome Fire impacts on microbial richness: Soil microbial richness was highly resistant to the Dome Fire for all three kingdoms. Low-intensity fires typically have subtle effects on microbes, impacting community connectivity and ecosystem services rather than richness (Aanderud et al., 2019; Soria et al., 2023; Vega-Cofre et al., 2023). Mojave National Preserve becomes primed for fire when high cool-and warm-season precipitation years are followed by a dry warm-season. These rainfall pulses and droughts also drive microbial dormancy and activation (G. Wang et al., 2014), consistent with our overall increase in biomass at year 3. However, an immediate post-fire dampening of species richness was not seen unlike high-intensity fires in shrublands and boreal systems (Holden et al., 2016; Pulido-Chavez et al., 2023). Frequent low-intensity fires can increase soil C and N, and this abundant substrate correlates with slightly increased bacterial richness in grasslands 1-2 months post-fire (Glassman et al., 2023), prairies sampled 1-month after fire (Mino et al., 2021) and pine forests sampled 1-2 years after fire (Fox et al., 2024). Here, bacterial richness was resistant in burned plots while unburned plots continued to drop at 3-years post-fire. Studies applying controlled burns to deserts varied from a similarly resistant bacterial community (Vega-Cofre et al., 2023) to a poorly resistant but highly resilient biocrust community (Aanderud et al., 2019). Archaeal richness did not differ post-fire, consistent with their greater resistance to fire even when fungal and bacterial richness decrease (Pérez-Valera et al., 2018). Fungal richness in drought-stricken grasslands showed greater resilience to fire than non-drought burned plots (Hacopian et al., 2024), suggesting that deserts, which have lower precipitation than grasslands, may also filter for highly resilient fungi. AMF richness was also greater in burned plots at 8-months and 3-years, decoupled from trends like plant richness and precipitation, perhaps due to fine-scale edaphic properties or plant dynamics not captured by our study. While no recent studies have comprehensively examined microbial richness after a desert wildfire, our results are aligned with low-intensity fires (Glassman et al., 2023; Vega-Cofre et al., 2023; Zhang et al., 2025) where bacterial and fungal richness rarely shift post-fire.

Dome Fire impacts on microbial biomass: Microbial biomass was resistant to fire, which is in line with biomass impacts of low intensity grassland and prescribed fires (Docherty et al., 2012; Pressler et al., 2019). Interestingly, archaeal and bacterial biomass increased in both plots by 3 years, after the influx of rain from tropical storm Hilary, but the abundant inorganic N may have contributed to the greater rise in burned compared to unburned plots. Prior to this rainfall, 16S copy numbers were in line with qPCR-based biomass estimates in desert biocrusts (Aanderud et al., 2019). Fungal biomass did not follow this pulse trend, rather it saw an increase in burned plots at 1-year post-fire potentially reacting to the widespread invasion of native annual and perennial herbs, a trend seen in other grasslands (Hennecke et al., 2025). AMF also show increases in spore abundance following some wildfires (Moura et al., 2022), reflecting a need for dormancy until vegetation recovers.

Direct versus indirect effects of Dome Fire on microbial composition: Fire had direct heat induced mortality effects on microbial community composition for all 3 kingdoms, and effects of fire were persistent with burned versus unburned microbial communities significantly differing for archaea for two and for bacteria and fungi at all timepoints. The limited archaeal response may be due to their lipid membrane structure and thick cell walls which makes them resistant to heat (Rothschild & Mancinelli, 2001). Archaeal and fungal communities appeared to trend towards divergence between burned and unburned rather than becoming more similar over the 3-year study, while the bacterial R^2^ did not seem to increase over time (Table S9). These divergences may be a result of fire increasing heterogeneity through pyrodiversity where a landscape does not burn uniformly (Hopkins et al., 2025), which is reflected in varied plant richness across the burn scar (Fig. S1). However, R^2^ values remained relatively small, consistent with grassland fires (Dove & Hart, 2017; Semenova-Nelsen et al., 2019) and much smaller than wildfires in forests and shrublands with high soil burn severity (Caiafa et al., 2023; Pulido-Chavez et al., 2023).

Drivers of microbial community composition were largely consistent with established global trends across differing kingdoms. Bacterial community composition was driven primarily by pH (R² = 0.38) and soil moisture (R² = 0.29), consistent with their overall resiliency to disturbance in dryland communities (Steven et al., 2021), well-known structuring by pH (Fierer & Jackson, 2006) and rapid response to precipitation in drylands (Blazewicz et al., 2014; Placella et al., 2012). Bacteria and archaea often show distinct community structuring, with one study showing nitrate driving bacterial but not archaeal communities (Johnson et al., 2017), inverse to our trend of nitrate correlating with both bacterial and archaeal communities while ammonium only linked to bacterial communities. Archaeal communities are often shaped by ammonium availability due to their activity as nitrifiers, oxidizing ammonium into nitrite during nitrification (Angel et al., 2010) and potentially driving the spikes in nitrate and nitrite found in select samples.

Fungal communities showed weaker correlations to environmental variables, driven by percent char (R² = 0.16) followed by pH (R² = 0.09) and plant richness (R² = 0.09), which was a significant driver of fungal composition but not bacterial or archaeal. The significant influence of plant richness on fungal communities was consistent with studies in temperate grasslands (Chen et al., 2017; Prober et al., 2015), but contrary to studies in desert systems that show soil, climate, and geographic distance drive fungal community composition in the unburned context (J. Wang et al., 2018; S. Wang et al., 2021). Archaeal communities were more similar to bacteria, however the influence of soil moisture (R² = 0.30) was slightly stronger than pH (R² = 0.27).

Studies of wetting desert soils show differential responses between domains, and although Archaea in general did not strongly respond to water, Ca. Nitrososphaera did increase in dominance within Archaea after wetting desert soil (Gao et al., 2021) and after a combined drying-wetting event (X.-B. Wang et al., 2022), suggesting species specific responses that may not be captured when assessing entire community dynamics. However, precipitation (considered as total from prior 1-month or 3-month) alone did not result in better models for prokaryotic richness or biomass, suggesting that further temporal dynamics beyond precipitation influenced prokaryotic communities. Meanwhile, soil moisture did not drive fungal community composition and yet fungal biomass was best modeled by the 3-month sum of prior precipitation, likely reflecting fungal associations with the plant community, as precipitation promotes plant growth, which in turn alters soil geochemistry and creates feedbacks that support fungal biomass. As bacterial and archaeal communities responded more directly to environmental conditions, fungal communities appeared to be impacted by char and the disappearance and reappearance of various plant species.

Impacts of Dome Fire on pyrophilous bacteria and archaea: Despite the relatively small changes in community composition, we identified a surprisingly high number of positive fire responding bacteria and archaea, including both known pyrophilous taxa and poorly described ASVs. More ASVs appeared respond positively to fire than were suppressed (Table S11), potentially allowing drought-tolerant, metabolically versatile microbes to increase in abundance post-fire. This is consistent with trends in forests immediately post-fire, in which bacteria are predominantly responding positively to fire and gradually even out over the span of a decade (Caiafa et al., 2023). After severe fires in shrublands and forests, Firmicutes typically increase whereas Acidobacteriota decrease (Enright et al., 2022; Nelson et al., 2024; Pérez-Valera et al., 2017; Pulido-Chavez et al., 2023; Rodríguez et al., 2018). In contrast, here, Firmicutes decreased and Acidobacteriota increased in overall relative abundance likely due to complex N-utilization of Pyrinomonadaceae members like Arenimicrobium and Blastocatella (Barriga et al., 2025; Wüst et al., 2016) (Fig. S5A). The Firmicutes genus Bacillus had greater relative abundance in unburned plots at every timepoint (Fig. S5B), however other pyrophilous Firmicutes like Tumebacillus and Paenibacillus were more abundant in burned plots, similar to other fires (Chungopast et al., 2023; Pulido-Chavez et al., 2023), potentially reflecting more post-fire specialization than the globally abundant Bacillus (Earl et al., 2008).

Other microbes responded according to their known metabolic activity. For example, nitrifier archaea like Ca. Nitrososphaera and Ca. Nitrocosmicus increased in burned plots, consistent with elevated ammonium (Han et al., 2024; Tourna et al., 2011), post-fire nitrogen pulses (Zhou et al., 2025), and their known drought tolerance (Stark & Firestone, 1995). Ca. Nitrososphaera oxidize ammonia into nitrite (Tourna et al., 2011), and are important keystone species in acidic soils for cycling N and aggregating soil (Banerjee et al., 2018; Y. Jiang et al., 2015). We detected previously identified pyrophilous Proteobacteria including Massilia (Caiafa et al., 2023; Fernández-González et al., 2023; Soria et al., 2023) and Noviherbasperillium (Pulido-Chavez et al., 2023; Woolet & Whitman, 2020). Their increase post-fire suggests conserved roles in post-fire soil recovery, even under the extreme drought and nutrient-limited conditions of the Mojave Desert. Bacteroidota members Adhaeribacter and Segetibacter, which we detected over 1-year post-fire, were positive fire responders associated with increased metabolic activity and total inorganic nitrogen in other burned soils as well (Adkins et al., 2022; Lucas-Borja et al., 2019). Several Actinobacteriota genera such as Asanoa, Crossiella, Cellulomonas, Rubrobacter, and Solirubrobacter likely benefit from similar shared traits that allow Actinobacteriota to increase in dominance post-fire (Whitman et al., 2019), such as thicker cell walls, a preference to higher pH, and a general stress tolerant physiology (Z.-M. Jiang et al., 2023; Norman et al., 2017). While we describe many taxa found increasing in other wildfire settings, several taxa remain unknown, highlighting the need for both trait-based and genomic investigations to understand post-fire succession in the desert.

Impact of Dome Fire on pyrophilous fungi: The desert mycobiome was dominated by Ascomycota, with a variety of genera from saprotrophic and endophytic classes such as Dothideomycetes, Eurotiomycetes, and Sordariomycetes showing positive responses to fire. However, unlike other wildfires, the Dome Fire caused an increase in the phylum Basidiomycota rather than a decrease (Fox et al., 2022; Pulido-Chavez et al., 2023), driven by mushroom-forming saprotrophic fungi such as Geastrum and Lepiota. Alternaria, a dark septate endophytic saprobe and potential plant pathogen in the class Dothideomycetes (Thomma, 2003), made up 5-19% of all fungal reads across our time series and was increased in burned plots at 8-months and 3-years (Fig. S8). Species in this genus may regulate their melanin content according to drought stress (Zuo et al., 2022), potentially increasing their fire resistance (Hopkins & Bennett, 2024). Another Dothideomycete genus, Preussia, showed increased abundance in burned plots from 2-weeks to 3-years post-fire which has been seen in controlled desert burns in China (Zhang et al., 2025). This genus forms hardy spores, which enhances their survival in drought conditions (Hacopian et al., 2024).

Despite small overall changes in community composition, a large number (143 ASVs) of fungal taxa showed positive log2fold increases in burned relative to unburned plots. Typically, there are as many negative as positive responders to fire (Caiafa et al., 2023; Revillini et al., 2022), however we only detected 33 fungal ASVs decreasing in burned plots, potentially due to adaptations for surviving in the arid desert pre-fire. Penicillium, a positive burn responder, is often dominant in both burned and unburned soils post-fire, and its role as a saprotroph makes it a key post-fire fungus that remains dominant in burned soil long-term, from 1 to 5 years post-fire (Whitman et al., 2019, 2025). Penicillium harbor a variety of genes for degrading hydrocarbons (Sari et al., 2025), and some species exhibit thermotolerance (van der Spuy, 1975). We identified four yeasts, one basidiomycete desert-dweller Naganishia (Canini et al., 2023; Lopez et al., 2024; Prober et al., 2015; Wei et al., 2022), and three ascomycetes Saitoella, Coniochaeta, and Aureobasidium, which showed positive differential abundance at 3-years in our study and in forest fires (Packard et al., 2023). Notably, Naganishia was more abundant in burned plots even 10 years after forest fires (Caiafa et al., 2023). Other yeast-like fungi were common extremophiles such as Neophaeococcomyces (Kurbessoian et al., 2024) and Knufia (Isola et al., 2022), which may have been pre-adapted to survive fire due to thermotolerance.

Through sequencing, we identified many fungi that are known to produce ascocarps post-fire. Consistent with our prediction, we saw an increase from members of Pyronemataceae such as Pseudotricharina, Scutellinia, and Spooneromyces, which are commonly found fruiting after high severity wildfires (Fox et al., 2022). Neurospora discreta, known for its heat-activated spores (Goddard, 1935), was observed growing abundantly on burned Y. jaegeriana bark but not Y. schidigera or Y. baccata. However, soil sequencing only identified aconidial species like N. terricola, highlighting different lifestyles in closely related species inhabiting the Mojave Desert.

## Conclusion

We performed the most comprehensive study of above and belowground direct and indirect effects of a desert wildfire to date and found that microbial biomass and richness were highly resistant in the face of roughly 80% mortality of Y. jaegeriana. The Dome Fire was a unique fire in that it was low intensity with low ash depth and char (0.37 ± 0.07 cm and 43.33% ± 4.37) but high severity with roughly 80% mortality of the dominant tree species. Although Y. jaegeriana mortality increased over time, there did not appear to be significant lag effects of microbes and in fact, certain groups like bacteria and AMF increased in richness in burned plots over time. The impacts on microbial communities were largely in line with low intensity grassland or prescribed fires (E. B. Allen et al., 2011; Brooks, 2002; Glassman et al., 2023), where microbes typically experience minimal changes in biomass or richness. Finally, surprisingly, despite subtle changes in the overall community composition, we identified a large number of taxa that positively responded to fire, and some of these were consistent with pyrophilous taxa that respond to high severity wildfires in shrublands and forests like Massilia and Noviherbaspirillum for bacteria, Ca. Nitrososphaera and Ca. Nitrocosmicus for archaea, and Penicillium, Coniochaeta, Aureobasidium, Alternaria, Naganishia, and Preussia for fungi, which responded in other desert fires. This study fills a knowledge gap and provides critical new insights into desert wildfire ecology by illustrating the high resistance and resilience of desert microbes in response to a low intensity but high severity wildfire.

**Table 1.** Model-estimated mean and standard error of soil chemical properties. Values in bold are significantly greater between pairwise treatments per timepoint, based on generalized mixed effects models.

| pH | Unburned | Burned | z-score | p value |
| --- | --- | --- | --- | --- |
| **Overall** | 7.05 ± 0.18 | **7.56 ± 0.14** | 3.13 | 0.001 |
| **17 Days** | 6.92 ± 0.06 | **7.63 ± 0.1** | -3.1 | 0.006 |
| 8 Months | 7.2 ± 0.12 | 7.58 ± 0.08 | -1.89 | 0.09 |
| **1 Year** | 6.68 ± 0.05 | **7.46 ± 0.09** | -3.5 | 0.012 |
| **3 Years** | 7.04 ± 0.03 | **7.49 ± 0.06** | -2.3 | 0.048 |
| Phosphate | Unburned | Burned | z-score | p value |
| Overall | 0.346 ± 0.1 | 0.46 ± 0.1 | 0.86 | 0.38 |
| 17 Days | 0.47 ± 0.39 | 0.36 ± 0.07 | -1.2 | 0.24 |
| 8 Months | 0.09 ± 0.03 | 0.15 ± 0.04 | -1.2 | 0.22 |
| 3 Years | 0.62 ± 0.24 | 0.51 ± 0.14 | 0.33 | 0.74 |
| Nitrite and Nitrate | Unburned | Burned | z-score | p value |
| **Overall** | 0.137 ± 0.04 | **0.324 ± 0.07** | 2.38 | 0.018 |
| 17 Days | 0.2 ± 0.09 | 0.32 ± 0.05 | -1.4 | 0.16 |
| 8 Months | 0.2 ± 0.01 | 0.24 ± 0.03 | -0.71 | 0.47 |
| **3 Years** | 0.01 ± 0.01 | **0.41 ± 0.23** | -2.1 | 0.04 |
| Ammonium | Unburned | Burned | z-score | p value |
| Overall | 0.3 ± 0.05 | 0.44 ± 0.05 | 1.6 | 0.11 |
| 17 Days | 0.6 ± 0.1 | 0.6 ± 0.08 | -0.043 | 0.96 |
| **8 Months** | 0.32 ± 0.06 | **0.51 ± 0.06** | -2.15 | 0.03 |
| **3 Years** | 0.13 ± 0.02 | **0.27 ± 0.03** | -3.3 | 0.001 |

## Notes

### Competing Interest Statement

The authors have declared no competing interest.

### Funding

- USDA-NIFA: #2022-67014-36675
- Shipley-Skinner Reserve - Riverside County Endowment

