## Supplemental Data for "Mojave Desert microbial communities show high resistance and resilience over three years despite widespread plant mortality following the Dome Fire"

**Contents**

Table S1. PCR thermocycler settings

Table S2. Plant richness and ash depth in burned and unburned plots

Table S3. Plant species identified across 3 field surveys at 17 days, 1 year, and 3 years

Table S4. Soil chemical properties across treatments and time

Table S5. Microbial richness across treatments and time

Table S6. Microbial biomass across treatments and time

Table S7. Model comparisons of bacterial richness and biomass with precipitation and time

Table S8. Model comparisons of fungal richness and biomass with precipitation and time

Table S9. Microbial Bray-Curtis dissimilarity between burned and unburned plots across time

Table S10. Variation partitioning of Bray-Curtis dissimilarity

Table S11. Positive differential abundance of prokaryotes in burned plots across time

Table S12. Negative differential abundance of prokaryotes in burned plots across time

Table S13. Positive differential abundance of fungi in burned plots across time

Table S14. Negative differential abundance of fungi in burned plots across time

Figure S1. Plant species richness by plot across time

Figure S2. Soil characteristics in burned and unburned plots across time

Figure S3. Fungal richness regressed against ash depth

Figure S4. Bacterial richness regressed against ash depth

Figure S5. Relative sequence abundance of archaeal and bacterial phyla and genera

Figure S6. Relative sequence abundance of fungal phyla and genera

Figure S7. Glomeromycotina relative sequence abundance

Figure S8. Fungal guild relative sequence abundance across time

Table S1. PCR thermocycler settings for PCR 1 (initial amplification), PCR 2 (attaching index and barcodes), 16S qPCR (archaeal and bacterial biomass estimation) and FungiQuant 18S qPCR (fungal biomass estimation).

| **Category** | **Initial Denaturation** | **Cycles** | **Denaturation** | **Annealing** | **Extension** | **Final Extension** |
| --- | --- | --- | --- | --- | --- | --- |
| **PCR1 (16S & ITS2)** | 94°C for 2 min | 30 | 94°C for 20 sec | 55°C for 20 sec | 68°C for 1 min | 68°C for 2 min |
| **PCR2 (DIP Primers)** | 94°C for 2 min | 10 | 94°C for 30 sec | 60°C for 30 sec | 72°C for 1 min | NA |
| **16S qPCR** | 94°C for 5 min | 40 | 94°C for 20 sec | 52°C for 30 sec | 72°C for 30 sec | NA |
| **18S FungiQuant qPCR** | 94°C for 5 min | 40 | 94°C for 20 sec | 50°C for 30 sec | 72°C for 30 sec | NA |

Table S2. Significant differences based on pairwise ANOVA between burned and unburned plots at each time point where plant richness and ash cover was recorded. Significant values bolded.

|  | **Plant Richness** | | | **Ash Cover at 17 Days** | | |
| --- | --- | --- | --- | --- | --- | --- |
|  | Pairwise ANOVA | | | Pairwise ANOVA | | |
|  | F_1,7_ | t value | p.value | F_1,34_ | t value | p.value |
| 17 Days | 49.2 | 11.1 | **0.001** | 144 | -12 | **8.9e-14** |
| 1 Year | 0.66 | -0.8 | 0.44 |  |  |  |
| 3 Years | 9.63 | -3.1 | **0.02** |  |  |  |

Table S3. Plant species appearing in our 3 field surveys in September 2020, 2021, and 2023 corresponding to 17 days, 1 year, and 3 years post-fire. Strata is represented by S = “shrub”, T = “Tree”, AH = “annual herb”, PH = “perennial herb”, H = “herb”, AG = “annual grass”, PG = “perennial grass”. Status is either N = “native”, N_R = “native, rare” or E = “exotic”. First Appearance indicates the sampling year in which a plant species was first observed. The Burned Plot Only column indicates if a plant species was only detected in a burned plot, with orange cells labeled “YES” and green cells labeled “NO”. Taxa not identified to genus or species are due to lack of identifiable material in the field.

| **Strata** | **Family** | **Scientific name** | **Status** | **First Appearance** | **Burned Plot Only** |
| --- | --- | --- | --- | --- | --- |
| S | Asteraceae | *Ambrosia salsola* | N | 2020 | NO |
| AG | Poaceae | *Bromus madritensis* | E | 2020 | NO |
| S | Rosaceae | *Coleogyne ramosissima* | N | 2020 | YES |
| S | Cactaceae | *Coryphantha alversonii* | N_R | 2020 | NO |
| S | Cactaceae | *Cylindropuntia acanthacarpa* | N | 2020 | NO |
| S | Cactaceae | *Cylindropuntia echinocarpa* | N | 2020 | NO |
| S | Cactaceae | *Cylindropuntia ramosisima* | N | 2020 | NO |
| PG | Poaceae | *Dasyochloa pulchella* | N | 2020 | NO |
| S | Ephedraceae | *Ephedra viridis* | N | 2020 | NO |
| S | Asteraceae | *Ericameria linearifolia* | N | 2020 | NO |
| S | Asteraceae | *Gutierrezia microcephala* | N | 2020 | YES |
| PH | Poaceae | *Hilaria rigida* | N | 2020 | NO |
| S | Solanaceae | *Lycium cooperi* | N | 2020 | NO |
| S | Lamiaceae | *Scutellaria mexicana* | N | 2020 | NO |
| S | Fabaceae | *Senna armata* | N | 2020 | NO |
| PH | Poaceae | *Stipa speciosa* | N | 2020 | NO |
| PH | Poaceae | *Stipa hymenoides* | N | 2020 | NO |
| S | Rutaceae | *Thamnosma montanta* | N | 2020 | NO |
| T | Agavaceae | *Yucca baccata* | N | 2020 | NO |
| T | Agavaceae | *Yucca jaegeriana* | N | 2020 | NO |
| T | Agavaceae | *Yucca schidigera* | N | 2020 | NO |
| S | Asteraceae | *Acamptopappus sphaerocephalus* | N | 2021 | NO |
| AH | Amaranthaceae | *Amaranthus fimbriatus* | N | 2021 | YES |
| PH | Fabaceae | *Astragalus sp.* | N | 2021 | YES |
| H | Nyctaginaceae | *Boerhavia sp.* | N | 2021 | YES |
| AG | Poaceae | *Bouteloua sp.* | N | 2021 | YES |
| AH | Polygonaceae | *Calyptridium monandrum* | N | 2021 | YES |
| S | Cactaceae | *Excobaria vivipara var. rosea* | N_R | 2021 | NO |
| S | Cactaceae | *Echinocereus engelmannii* | N | 2021 | NO |
| S | Cactaceae | *Echinocereus mojavensis* | N | 2021 | NO |
| S | Ephedraceae | *Ephedra nevadensis* | N | 2021 | NO |
| AH | Geraniaceae | *Erodium cicutarium* | E | 2021 | YES |
| AH | Euphorbiaceae | *Euphorbia albomarginata* | N | 2021 | YES |
| AH | Euphorbiaceae | *Euphorbia sp.* | N | 2021 | YES |
| S | Poaceae | *Hilaria jamesii* | N | 2021 | YES |
| S | Kramariaceae | *Krameria sp.* | N | 2021 | YES |
| S | Solanaceae | *Lycium andersonii* | N | 2021 | NO |
| S | Cactaceae | *Mammillaria tetrancistra* | N | 2021 | NO |
| S | Oleaceae | *Menodora spinescens* | N | 2021 | YES |
| PH | Nyctaginaceae | *Mirabilis laevis* | N | 2021 | YES |
| H | Asteraceae | *Pectis papposa* | N | 2021 | YES |
| AH | Poaceae | *Poaceae sp. (Annual)* | N | 2021 | YES |
| H | Malvaceae | *Sphaeralcea ambigua* | N | 2021 | YES |
| PH | Malvaceae | *Sphaeralcea sp.* | N | 2021 | YES |
| PH | Poaceae | *Stipa sp.* | N | 2021 | YES |
| AH | Unknown | Annual herb/forb sp. | N | 2021 | YES |
| S | Asteraceae | *Ambrosia dumosa* | N | 2023 | NO |
| AG | Poaceae | *Aristida purpurea* | N | 2023 | YES |
| PG | Poaceae | *Bouteloua eriopoda* | N | 2023 | YES |
| S | Asteraceae | *Ericameria cooperi* | N | 2023 | NO |
| PH | Nyctaginaceae | *Mirabilis multiflora* | N | 2023 | YES |
| S | Poaceae | *Muhlenbergia porteri* | N | 2023 | NO |
| PH | Malvaceae | *Sphaeralcea rusbyi var. eremicola* | N_R | 2023 | YES |
| PH | Poaceae | *Poaceae sp (Perennial)* | N | 2023 | NO |

Table S4. Generalized mixed effect model statistics for soil chemical properties of pH, phosphate (PO_4_^3-^), ammonium (NH_4_^+^), and nitrate (NO_3_^-^ plus nitrite (NO_2_-) by fire, time, and time by fire interactions. Timepoints indicate comparisons across the unburned reference level. Time and fire interactions are listed below the overall treatment effect “Burned”. Marginal R^2^ reports model effect size after removing random effects. Significant values bolded.

|  | **pH** | | | | | | **PO_4_^3-^** | | | | | |
| --- | --- | --- | --- | --- | --- | --- | --- | --- | --- | --- | --- | --- |
|  | Marginal R^2^ =0.33 | | | | | | Marginal R^2^ =0.29 | | | | | |
|  | estimate | | z value | p.value | | | estimate | | | z value | | p.value |
| (Intercept) | 1.93 | | 87.30 | **<0.01** | | | -1.91 | | | -4.52 | | **<0.01** |
| Burned | 0.10 | | 3.54 | **0.01** | | | 0.59 | | | 1.18 | | 0.24 |
| 8 Months | 0.04 | | 2.99 | **<0.01** | | | -0.56 | | | -1.20 | | 0.23 |
| 3 Years | 0.02 | | 1.32 | 0.19 | | | 1.44 | | | 3.07 | | **<0.01** |
| 8 Months:Burned | -0.04 | | -2.89 | **0.01** | | | -0.01 | | | -0.02 | | 0.99 |
| 3 Years:Burned | -0.03 | | -2.19 | **0.03** | | | -0.75 | | | -1.31 | | 0.19 |
|  | **NH_4_^+^** | | | | | | **NO_3_^-^ and NO_2_^-^** | | | | | |
|  | Marginal R^2^ =0.61 | | | | | | Marginal R^2^ =0.46 | | | | | |
|  | estimate | | z value | p.value | | | estimate | | | z value | | p.value |
| (Intercept) | -0.51 | | -2.90 | **<0.01** | | | -1.70 | | | -2.33 | | **0.0002** |
| Burned | 0.01 | | 0.04 | 0.97 | | | 1.29 | | | 1.40 | | 0.36 |
| 8 Months | -0.63 | | -4.72 | **<0.01** | | | 0.47 | | | 0.85 | | 0.99 |
| 3 Years | -1.51 | | -11.26 | **<0.01** | | | -6.63 | | | -10.89 | | **<0.001** |
| 8 Months:Burned | 0.45 | | 2.76 | **0.01** | | | -0.64 | | | -0.94 | | 0.69 |
| 3 Years:Burned | 0.70 | | 4.29 | **<0.01** | | | 1.20 | | | 0.97 | | **<0.001** |
| Table S5. Generalized mixed effect model statistics for amplicon sequence variant (ASV) richness of bacteria, fungi, archaea, and arbuscular mycorrhizal fungi (AMF) by fire, time, and fire by time interactions. Timepoints indicate comparisons across the unburned reference level. Time and fire interactions are listed below the overall treatment effect “Burned”. Marginal R^2^ reports model effect size after removing random effects. | | | | | | | | | | | | |
|  | **Bacteria** | | | | | **Fungi** | | | | | | |
| **ASV Richness** |  | Marginal R^2^ = 0.176 | | |  |  | | Marginal R^2^ = 0.194 | | | | |
|  | estimate | | z value | p.value | | estimate | | | z value | | p.value | |
| (Intercept) | 6.37 | | 54.83 | **<0.01** | | 168.75 | | | 10.74 | | **<0.01** | |
| 1 Month | 0.11 | | 0.55 | 0.59 | | -24.78 | | | -1.01 | | 0.32 | |
| 8 Months | -0.41 | | -3.02 | **0.003** | | -33.56 | | | -1.87 | | 0.06 | |
| 1 Year | -0.29 | | -2.12 | **0.036** | | -17.17 | | | -1.04 | | 0.30 | |
| 3 Years | -0.54 | | -3.97 | 0.0001 | | 24.21 | | | 1.43 | | 0.16 | |
| Burned | -0.19 | | -1.34 | 0.19 | | 2.50 | | | 0.13 | | 0.90 | |
| 1 Month:Burned | 0.11 | | 0.51 | 0.61 | | -5.22 | | | -0.19 | | 0.85 | |
| 8 Months:Burned | 0.29 | | 1.77 | 0.08 | | 18.62 | | | 0.86 | | 0.39 | |
| 1 Year:Burned | 0.20 | | 1.20 | 0.23 | | 6.33 | | | 0.31 | | 0.76 | |
| 3 Years:Burned | 0.51 | | 3.07 | **0.0025** | | 10.54 | | | 0.51 | | 0.61 | |
|  | **Archaea** | | | | | **AMF** | | | | | | |
| **ASV Richness** | Marginal R^2^ =0.11 | | | | | Marginal R^2^ = 0.31 | | | | | | |
|  | estimate | | z value | p.value | | estimate | | | z value | | p.value | |
| (Intercept) | 2.92 | | 28.38 | <0.01 | | 1.79 | | | 6.62 | | **<0.01** | |
| 1 MONTH | 0.06 | | 0.41 | 0.69 | | 0.38 | | | 0.80 | | 0.43 | |
| 8 MONTHS | -0.23 | | -2.10 | 0.04 | | -0.71 | | | -2.32 | | **0.02** | |
| 1 Year | -0.17 | | -1.54 | 0.12 | | -0.34 | | | -1.10 | | 0.27 | |
| 3 Years | -0.26 | | -2.27 | 0.02 | | 0.16 | | | 0.56 | | 0.58 | |
| Burned | -0.01 | | -0.04 | 0.96 | | 0.45 | | | 1.33 | | 0.18 | |
| 1 Month:Burned | 0.01 | | 0.03 | 0.98 | | -1.12 | | | -2.15 | | **0.03** | |
| 8 Months:Burned | 0.14 | | 1.04 | 0.30 | | 0.26 | | | 0.69 | | 0.49 | |
| 1 Year:Burned | 0.01 | | 0.07 | 0.94 | | -0.27 | | | -0.72 | | 0.47 | |
| 3 Years:Burned | 0.17 | | 1.23 | 0.22 | | 0.44 | | | 1.22 | | 0.22 | |

Table S6. Generalized mixed effect model statistics for copy numbers of 16S and 18S. Timepoints indicate comparisons across the unburned reference level. Time and fire interactions are listed below the overall treatment effect “Burned”. Marginal R^2^ reports model effect size after removing random effects.

|  | **Bacteria (16S)** | | | | | **Fungi (18S)** | | | | |
| --- | --- | --- | --- | --- | --- | --- | --- | --- | --- | --- |
| **Biomass**  (qPCR Copy Number) |  | Marginal R^2^: 0.924 | | |  | |  | Marginal R^2^: 0.353 | | |
|  | estimate | | z value | p value | | estimate | | | z value | p value |
| (Intercept) | 18.38 | | 155.77 | **<0.01** | | 14.86 | | | 94.59 | **<0.01** |
| 1 Month | -0.22 | | -1.09 | 0.28 | | 0.18 | | | 0.36 | 0.72 |
| 8 Months | -0.64 | | -5.32 | **<0.01** | | -0.22 | | | -1.19 | 0.23 |
| 1 Year | -0.94 | | -7.81 | **<0.01** | | -1.12 | | | -3.06 | **<0.01** |
| 3 Years | 1.97 | | 15.61 | **<0.01** | | -1.29 | | | -3.05 | **<0.01** |
| Burned | 0.15 | | 1.06 | 0.30 | | -0.03 | | | -0.15 | 0.88 |
| 1 Month:Burned | -0.14 | | -0.66 | 0.51 | | -0.39 | | | -0.70 | 0.48 |
| 8 Months:Burned | -0.03 | | -0.21 | 0.83 | | 0.16 | | | 0.73 | 0.46 |
| 1 Year:Burned | 0.08 | | 0.51 | 0.61 | | 0.86 | | | 2.18 | **0.03** |
| 3 Years:Burned | 0.34 | | 2.17 | **0.03** | | 0.14 | | | 0.28 | 0.78 |

Table S7. Bacterial model comparisons of time as categorical (TimePoint), numerical (Days Post-fire), and precipitation as a sum of 3-months prior, 1 month-prior, and soil moisture. Limiting timepoints to include plant richness did not improve model fidelity. Each model comparison for biomass and richness had the variable in the top row as a fixed effect and Plot as a random effect. Models are arranged from highest to lowest Akaike Information Criteron (AIC). Delta AIC indicates the difference between the null model AIC and each model’s own AIC. Chi squared indicates the improvement of the model fit compared to the null model. In the case of < 3 AIC difference, the greater Chi Squared value is compared for best fit. Bolded columns indicate the best fit model as determined by comparison of Chi squared and AIC for biomass and richness.

| **Biomass** | Null model | Soil Moisture | Precipitation (3 month sum) | Days Post-fire | Precipitation (1 month sum) | **TimePoint** |
| --- | --- | --- | --- | --- | --- | --- |
| AIC | 6450.5 | 6400 | 6332.5 | 6227.6 | 6192.3 | **5970.8** |
| Delta AIC | 0 | 50.5 | 118 | 222.9 | 258.2 | **479.7** |
| Chi Squared |  | 56.51 | 67.54 | 0 | 140.14 | **268.79** |
| p value | NA | 3.28E-12 | <2.2e-16 | 1 | <2.2e-16 | **<2.2e-16** |
| **Richness** | Null model | Precipitation (1 month sum) | Precipitation (3 month sum) | Days Post-fire | Soil Moisture | **TimePoint** |
| AIC | 152.4 | 149.6 | 145.1 | 144.1 | 142.9 | **133.9** |
| Delta AIC | 0 | 2.9 | 7.3 | 8.3 | 9.6 | **18.5** |
| Chi Squared |  | 0 | 11.31 | 0 | 8.7 | **20.09** |
| p value | NA | n.s. | 0.0035 | n.s. | 0.0032 | **0.0027** |

Table S8. Fungal model comparisons of time as categorical (TimePoint), numerical (Days Post-fire), and precipitation as a sum of 3-months prior, 1 month-prior, and soil moisture. Limiting timepoints to include plant richness did not improve model fidelity. Each model comparison for biomass and richness had the variable in the top row as a fixed effect and Plot as a random effect. Models are arranged from highest to lowest Akaike Information Criteron (AIC). Delta AIC indicates the difference between the null model AIC and each model’s own AIC. Chi squared indicates the improvement of the model fit compared to the null model. In the case of < 3 AIC difference, the greater Chi Squared value is compared for best fit. Bolded columns indicate the best fit model as determined by comparison of Chi squared and AIC for biomass and richness.

| **Biomass** | | Null model | Soil Moisture | **Precipitation (3 month sum)** | TimePoint | Days Post-fire | Precipitation (1 month sum) |
| --- | --- | --- | --- | --- | --- | --- | --- |
|  | AIC | 5329 | 5324.2 | **5292.5** | 5290.9 | 5290.1 | 5289.6 |
|  | Delta AIC | 0 | 4.8 | **36.5** | 38.1 | 38.9 | 39.4 |
|  | Chi Squared |  | 10.75 | **31.73** | 11.14 | 0 | 2.85 |
|  | p value | NA | 0.013 | **<2e-16** | 0.084 | 1 | <2e-16 |
| **Richness** | | Null model | Precipitation (3 month sum) | Soil Moisture | Days Post-fire | Precipitation (1 month sum) | **TimePoint** |
|  | AIC | 1732.5 | 1723.1 | 1722.9 | 1710.9 | 1708.3 | **1707.6** |
|  | Delta AIC | 0 | 9.4 | 9.6 | 21.6 | 24.2 | **24.9** |
|  | Chi Squared |  | 0 | 0 | 0 | 14.74 | **27.32** |
|  | p value | NA | n.s. | n.s. | n.s. | n.s. | **0.0001** |

Table S9. Community composition differences between burned and unburned plots in Archaea, Bacteria, and Fungi. PERMANOVA and Beta-dispersion are compared at each timepoint and shown in bold when significant. A significant PERMANOVA indicates differing centroid means and suggests differing community composition. A significant beta-dispersion indicates differing distance to centroid and suggests differing levels of heterogeneity. A combination of significant PERMANOVA and betadispersion suggests differing communities due to differences in both composition and heterogeneity.

| Domain | Time | Burn R^2^ | | PERM-ANOVA F | | p-value | |  | Betadisper F | | | p-value |
| --- | --- | --- | --- | --- | --- | --- | --- | --- | --- | --- | --- | --- |
| Archaea | 17 Days | 0.058 | | 1.97 | | 0.065 | |  | 2.84 | | | 0.10 |
|  | 1 Month | 0.062 | | 1.74 | | 0.101 | |  | 9.51 | | | **0.004** |
|  | 8 Months | 0.069 | | 2.55 | | **0.021** | |  | 3.18 | | | 0.083 |
|  | 1-Year | 0.05 | | 1.78 | | 0.072 | |  | 4.05 | | | 0.052 |
|  | 3-Years | 0.073 | | 2.37 | | **0.014** | |  | 1.57 | | | 0.2185 |
| Bacteria | 17 Days | 0.056 | | 2.01 | | **0.001** | |  | 9.06 | | | **0.004** |
|  | 1 Month | 0.052 | | 1.42 | | **0.032** | |  | 11.95 | | | **0.0018** |
|  | 8 Months | 0.045 | | 1.6 | | **0.007** | |  | 0.49 | | | 0.48 |
|  | 1-Year | 0.063 | | 2.32 | | **0.001** | |  | 3.44 | | | 0.072 |
|  | 3-Years | 0.047 | | 1.69 | | **0.001** | |  | 2.51 | | | 0.12 |
| Fungi | 17 Days | 0.068 | | 2.48 | | **0.001** | |  | 12.09 | | | **0.0014** |
|  | 1 Month | 0.059 | | 1.65 | | **0.003** | |  | 18.76 | | | **0.00019** |
|  | 8 Months | 0.071 | | 2.32 | | **0.001** | |  | 0.89 | | | 0.35 |
|  | 1-Year | 0.063 | | 2.24 | | **0.001** | |  | 3.62 | | | 0.065 |
|  | 3-Years | 0.085 | | 3.07 | | **0.001** | |  | 0.41 | | | 0.52 |

Table S10. Variation partitioning of Bray–Curtis dissimilarity between burned and unburned communities per timepoint and per kingdom. The analysis separates the variance in community composition explained by environmental variables (only those significant in Figure 4), spatial distance based on subplot coordinates, their shared effects, and the unexplained remainder. Fractions represent the proportion of variation attributed to each component, with negative fractions occurring due to statistical adjustments and interpreted as zero.

| **Bacteria** |  | | |
| --- | --- | --- | --- |
| Fraction | 17 Days | 8 Months | 3 Years |
| Pure Environment | -0.024 | 0.133 | 0.077 |
| Pure Space | 0.251 | 0.091 | 0.123 |
| Shared (Env ∩ Space) | 0.044 | 0.061 | 0.042 |
| Unexplained | 0.729 | 0.715 | 0.758 |
| **Archaea** |  | | |
| Fraction | 17 Days | 8 Months | 3 Years |
| Pure Environment | -0.061 | 0.068 | 0.081 |
| Pure Space | 0.183 | 0.086 | -0.024 |
| Shared (Env ∩ Space) | 0 | 0.04 | -0.025 |
| Unexplained | 0.879 | 0.806 | 0.968 |

| **Fungi** |  | | |
| --- | --- | --- | --- |
| Fraction | 17 Days | 1 Year | 3 Years |
| Pure Environment | 0.031 | 0.1 | 0.065 |
| Pure Space | 0.208 | 0.278 | 0.145 |
| Shared (Env ∩ Space) | 0.099 | 0.081 | 0.034 |
| Unexplained | 0.662 | 0.54 | 0.756 |

Table S11. Prokaryotic genera showing positive differential abundance in burned plots (increase) when compared to unburned plots, sorted by timepoint and kingdom. All rows showed significant increases in burned plots based on the adjusted p value (padj < 0.01) after a Benjamini–Hochberg correction in DESEQ2.

| log2Fold Change | padj | Kingdom | Phylum | Genus | Species | Timepoint |
| --- | --- | --- | --- | --- | --- | --- |
| 22.08 | 2E-11 | Bacteria | Firmicutes | *Tumebacillus* | *NA* | 17 Days |
| 22.24 | 3E-11 | Bacteria | Actinobacteriota | *Crossiella* | uncultured bacterium | 8 Months |
| 22.10 | 1E-09 | Bacteria | Acidobacteriota | Vicinamibacteraceae | *NA* | 8 Months |
| 22.45 | 2E-20 | Bacteria | Actinobacteriota | *NA* | *NA* | 1 Year |
| 22.00 | 1E-12 | Bacteria | Proteobacteria | *NA* | *NA* | 1 Year |
| 21.99 | 1E-12 | Bacteria | Actinobacteriota | *Asanoa* | *NA* | 1 Year |
| 21.97 | 1E-12 | Bacteria | Acidobacteriota | Vicinamibacteraceae | *NA* | 1 Year |
| 21.96 | 2E-10 | Bacteria | Proteobacteria | *NA* | *NA* | 1 Year |
| 21.76 | 1E-12 | Bacteria | Verrucomicrobiota | *Candidatus Udaeobacter* | *NA* | 1 Year |
| 21.61 | 4E-10 | Bacteria | Myxococcota | *bacteriap25* | *NA* | 1 Year |
| 21.58 | 4E-10 | Bacteria | Acidobacteriota | *uncultured* | uncultured bacterium | 1 Year |
| 21.38 | 5E-10 | Bacteria | Proteobacteria | *NA* | *NA* | 1 Year |
| 21.20 | 7E-10 | Bacteria | Actinobacteriota | *Rubrobacter* | uncultured actinobacterium | 1 Year |
| 20.84 | 2E-09 | Bacteria | Proteobacteria | *NA* | *NA* | 1 Year |
| 20.45 | 3E-09 | Bacteria | Proteobacteria | *Noviherbaspirillum* | *NA* | 1 Year |
| 7.23 | 5E-05 | Bacteria | Bacteroidota | *Arcticibacter* | uncultured bacterium | 1 Year |
| 4.34 | 3E-05 | Bacteria | Firmicutes | *NA* | *NA* | 1 Year |
| 3.18 | 3E-03 | Bacteria | Proteobacteria | *Massilia* | *NA* | 1 Year |
| 2.19 | 2E-08 | Bacteria | Proteobacteria | *Massilia* | *NA* | 1 Year |
| 21.88 | 2E-11 | Archaea | Crenarchaeota | *Candidatus Nitrocosmicus* | *NA* | 3 Years |
| 21.09 | 5E-09 | Archaea | Crenarchaeota | *Candidatus Nitrososphaera* | *NA* | 3 Years |
| 21.97 | 1E-09 | Bacteria | Actinobacteriota | *Solirubrobacter* | *NA* | 3 Years |
| 22.42 | 7E-15 | Bacteria | Actinobacteriota | *Rubrobacter* | uncultured *Rubrobacter* | 3 Years |
| 21.34 | 3E-09 | Bacteria | Firmicutes | *NA* | *NA* | 3 Years |
| 21.29 | 3E-09 | Bacteria | Bacteroidota | *Segetibacter* | *NA* | 3 Years |
| 20.57 | 1E-08 | Bacteria | Actinobacteriota | *Solirubrobacter* | *NA* | 3 Years |
| 7.15 | 2E-04 | Bacteria | Actinobacteriota | *Cellulomonas* | *NA* | 3 Years |

Table S12. Prokaryotic genera showing negative differential abundance in burned plots (decrease) when compared to unburned plots, sorted by timepoint. All rows showed significant decreases in burned plots based on the adjusted p value (padj < 0.01) after a Benjamini–Hochberg correction in DESEQ2.

| log2Fold Change | padj | Kingdom | Phylum | Genus | Species | Timepoint |
| --- | --- | --- | --- | --- | --- | --- |
| -22.95 | 2E-10 | Bacteria | Actinobacteriota | *Gaiella* | *NA* | 17 Days |
| -22.95 | 2E-10 | Bacteria | Actinobacteriota | uncultured | uncultured bacterium | 17 Days |
| -23.40 | 5E-11 | Bacteria | Myxococcota | *bacteriap25* | metagenome | 8 Months |
| -23.54 | 5E-11 | Bacteria | Actinobacteriota | *NA* | *NA* | 8 Months |
| -24.24 | 1E-12 | Bacteria | Cyanobacteria | *uncultured* | *Scytonema hyalinum* | 1 Year |
| -24.66 | 6E-13 | Bacteria | Actinobacteriota | *Rubrobacter* | uncultured *Rubrobacter* | 1 Year |
| -24.47 | 3E-12 | Bacteria | Firmicutes | *Psychrobacillus* | *NA* | 3 Years |

Table S13. Fungal genera showing positive differential abundance in burned plots (increase) when compared to unburned plots. All rows showed significant increases in burned plots based on the adjusted p value (padj < 0.01) after a Benjamini–Hochberg correction in DESEQ2.

| log2Fold Change | padj | Kingdom | Phylum | Genus | Species | Timepoint |
| --- | --- | --- | --- | --- | --- | --- |
| 24.12 | 4E-13 | Fungi | Ascomycota | Pezizaceae inc. sed. | Pezizaceae sp | 17 Days |
| 24.09 | 4E-13 | Fungi | Basidiomycota | *Geastrum* | *Geastrum sp* | 17 Days |
| 23.68 | 2E-25 | Fungi | Ascomycota | *NA* | *NA* | 17 Days |
| 23.59 | 4E-13 | Fungi | Ascomycota | *Knufia* | *NA* | 17 Days |
| 23.59 | 1E-12 | Fungi | Ascomycota | *Ascobolus* | *NA* | 17 Days |
| 23.17 | 1E-23 | Fungi | Ascomycota | *Preussia* | *Preussia fleischhakii* | 17 Days |
| 22.86 | 5E-12 | Fungi | Ascomycota | *Humicola* | *NA* | 17 Days |
| 22.82 | 1E-16 | Fungi | Ascomycota | *Gibberella* | *Gibberella avenacea* | 17 Days |
| 22.79 | 9E-21 | Fungi | Ascomycota | *Aureobasidium* | *NA* | 17 Days |
| 22.63 | 8E-12 | Fungi | Ascomycota | *Preussia* | *Preussia sp* | 17 Days |
| 22.58 | 8E-12 | Fungi | Ascomycota | *Talaromyces* | *NA* | 17 Days |
| 22.38 | 1E-11 | Fungi | Ascomycota | *Acrophialophora* | *Acrophialophora sp* | 17 Days |
| 22.33 | 3E-17 | Fungi | Ascomycota | *Articulospora* | *NA* | 17 Days |
| 22.20 | 7E-12 | Fungi | Ascomycota | *Acrophialophora* | *Acrophialophora sp* | 17 Days |
| 22.02 | 3E-11 | Fungi | Ascomycota | *NA* | *NA* | 17 Days |
| 21.83 | 4E-11 | Fungi | Ascomycota | *Coniochaeta* | *NA* | 17 Days |
| 21.71 | 5E-11 | Fungi | Ascomycota | Chaetothyriales inc. sed. | Chaetothyriales sp | 17 Days |
| 21.67 | 5E-11 | Fungi | Ascomycota | *Aspergillus* | *NA* | 17 Days |
| 21.65 | 5E-11 | Fungi | Ascomycota | *Pseudotricharina* | *Pseudotricharina sp* | 17 Days |
| 21.60 | 6E-11 | Fungi | Ascomycota | *NA* | *NA* | 17 Days |
| 21.29 | 1E-10 | Fungi | Ascomycota | *NA* | *NA* | 17 Days |
| 20.91 | 2E-10 | Fungi | Ascomycota | *Chaetomium* | *NA* | 17 Days |
| 20.16 | 3E-11 | Fungi | Ascomycota | *Hazslinszkyomyces* | *Hazslinszkyomyces lycii* | 17 Days |
| 8.91 | 1E-05 | Fungi | Ascomycota | *Fusarium* | *NA* | 17 Days |
| 7.23 | 7E-03 | Fungi | Ascomycota | *Knufia* | *Knufia sp* | 17 Days |
| 6.34 | 9E-03 | Fungi | Ascomycota | *NA* | *NA* | 17 Days |
| 23.68 | 6E-11 | Fungi | Ascomycota | *NA* | *NA* | 1 Month |
| 22.73 | 4E-08 | Fungi | Basidiomycota | *Geastrum* | *Geastrum sp* | 1 Month |
| 22.45 | 4E-06 | Fungi | Ascomycota | *NA* | *NA* | 1 Month |
| 22.27 | 2E-09 | Fungi | Ascomycota | *Neophaeococcomyces* | *Neophaeococcomyces sp* | 1 Month |
| 21.90 | 4E-10 | Fungi | Ascomycota | *Neophaeococcomyces* | *Neophaeococcomyces sp* | 1 Month |
| 21.86 | 9E-11 | Fungi | Ascomycota | *NA* | *NA* | 1 Month |
| 21.68 | 4E-08 | Fungi | Ascomycota | *Knufia* | *NA* | 1 Month |
| 21.63 | 1E-05 | Fungi | Ascomycota | *Acrophialophora* | *Acrophialophora sp* | 1 Month |
| 21.15 | 2E-08 | Fungi | Ascomycota | *NA* | *NA* | 1 Month |
| 21.01 | 1E-05 | Fungi | Ascomycota | *NA* | *NA* | 1 Month |
| 20.85 | 2E-05 | Fungi | Ascomycota | *Knufia* | *NA* | 1 Month |
| 20.77 | 1E-10 | Fungi | Ascomycota | *Sporormia* | *Sporormia sp* | 1 Month |
| 20.70 | 2E-05 | Fungi | Ascomycota | *Gibberella* | *NA* | 1 Month |
| 20.62 | 1E-05 | Fungi | Ascomycota | Didymellaceae inc. sed. | Didymellaceae sp | 1 Month |
| 20.41 | 1E-05 | Fungi | Ascomycota | *NA* | *NA* | 1 Month |
| 20.39 | 3E-05 | Fungi | Ascomycota | *Chaetosphaeronema* | *Chaetosphaeronema sp* | 1 Month |
| 20.16 | 4E-05 | Fungi | Ascomycota | *Pyrenochaeta* | *Pyrenochaeta sp* | 1 Month |
| 19.86 | 5E-05 | Fungi | Ascomycota | *NA* | *NA* | 1 Month |
| 18.80 | 2E-04 | Fungi | Ascomycota | *NA* | *NA* | 1 Month |
| 9.41 | 2E-04 | Fungi | Ascomycota | *Chaetosphaeronema* | *Chaetosphaeronema sp* | 1 Month |
| 23.97 | 3E-28 | Fungi | Ascomycota | *Neophaeococcomyces* | *Neophaeococcomyces sp* | 8 Months |
| 23.96 | 6E-13 | Fungi | Ascomycota | *NA* | *NA* | 8 Months |
| 22.94 | 8E-13 | Fungi | Ascomycota | *Darksidea* | *Darksidea alpha* | 8 Months |
| 22.61 | 3E-17 | Fungi | Ascomycota | *Trichoderma* | *NA* | 8 Months |
| 22.52 | 2E-11 | Fungi | Ascomycota | *NA* | *NA* | 8 Months |
| 22.45 | 5E-20 | Fungi | Ascomycota | *NA* | *NA* | 8 Months |
| 22.25 | 6E-13 | Fungi | Ascomycota | *Podospora* | *Podospora sp* | 8 Months |
| 22.16 | 3E-11 | Fungi | Ascomycota | *Acrophialophora* | *Acrophialophora sp* | 8 Months |
| 21.94 | 5E-11 | Fungi | Ascomycota | *Podospora* | *Podospora intestinacea* | 8 Months |
| 21.87 | 5E-11 | Fungi | Ascomycota | *NA* | *NA* | 8 Months |
| 21.67 | 8E-11 | Fungi | Ascomycota | *Preussia* | *Preussia terricola* | 8 Months |
| 21.59 | 1E-12 | Fungi | Ascomycota | *NA* | *NA* | 8 Months |
| 21.50 | 1E-10 | Fungi | Ascomycota | *Chaetosphaeronema* | *Chaetosphaeronema sp* | 8 Months |
| 21.47 | 1E-10 | Fungi | Ascomycota | *Alternaria* | *NA* | 8 Months |
| 21.45 | 1E-10 | Fungi | Ascomycota | *Clarireedia* | *Clarireedia bennettii* | 8 Months |
| 21.40 | 1E-10 | Fungi | Ascomycota | *NA* | *NA* | 8 Months |
| 21.24 | 2E-10 | Fungi | Ascomycota | *Pseudotricharina* | *Pseudotricharina sp* | 8 Months |
| 21.22 | 2E-10 | Fungi | Ascomycota | *NA* | *NA* | 8 Months |
| 20.88 | 3E-10 | Fungi | Ascomycota | *Preussia* | *NA* | 8 Months |
| 8.71 | 4E-03 | Fungi | Ascomycota | *Preussia* | *Preussia polymorpha* | 8 Months |
| 6.51 | 2E-11 | Fungi | Ascomycota | *NA* | *NA* | 8 Months |
| 24.86 | 8E-14 | Fungi | Ascomycota | *NA* | *NA* | 1 Year |
| 24.78 | 4E-20 | Fungi | Ascomycota | *Gibberella* | *NA* | 1 Year |
| 24.26 | 2E-13 | Fungi | Ascomycota | *Pseudotricharina* | *Pseudotricharina sp* | 1 Year |
| 23.77 | 4E-16 | Fungi | Ascomycota | *Ascobolus* | *NA* | 1 Year |
| 23.46 | 3E-13 | Fungi | Ascomycota | *Saitoella* | *Saitoella complicata* | 1 Year |
| 23.39 | 4E-24 | Fungi | Basidiomycota | *Naganishia* | *Naganishia albida* | 1 Year |
| 23.28 | 2E-12 | Fungi | Ascomycota | *Pyrenochaeta* | *Pyrenochaeta sp* | 1 Year |
| 23.13 | 2E-12 | Fungi | Ascomycota | *Spooneromyces* | *Spooneromyces laeticolor* | 1 Year |
| 23.13 | 3E-13 | Fungi | Ascomycota | *Darksidea* | *Darksidea alpha* | 1 Year |
| 22.91 | 4E-12 | Fungi | Ascomycota | *NA* | *NA* | 1 Year |
| 22.78 | 5E-12 | Fungi | Ascomycota | *Knufia* | *NA* | 1 Year |
| 22.66 | 1E-13 | Fungi | Ascomycota | *NA* | *NA* | 1 Year |
| 22.58 | 7E-12 | Fungi | Ascomycota | *Preussia* | *NA* | 1 Year |
| 22.44 | 1E-11 | Fungi | Basidiomycota | *Naganishia* | *NA* | 1 Year |
| 22.38 | 1E-13 | Fungi | Ascomycota | *NA* | *NA* | 1 Year |
| 22.37 | 1E-11 | Fungi | Ascomycota | Didymellaceae inc. sed. | Didymellaceae sp | 1 Year |
| 22.35 | 1E-11 | Fungi | Ascomycota | *NA* | *NA* | 1 Year |
| 22.22 | 1E-11 | Fungi | Ascomycota | *Mycocalicium* | *Mycocalicium victoriae* | 1 Year |
| 21.88 | 3E-11 | Fungi | Ascomycota | *Knufia* | *NA* | 1 Year |
| 21.73 | 4E-11 | Fungi | Ascomycota | *NA* | *NA* | 1 Year |
| 21.61 | 5E-11 | Fungi | Ascomycota | *Iodophanus* | *NA* | 1 Year |
| 21.56 | 5E-11 | Fungi | Ascomycota | *NA* | *NA* | 1 Year |
| 21.34 | 8E-11 | Fungi | Basidiomycota | *Naganishia* | *Naganishia albida* | 1 Year |
| 21.33 | 8E-11 | Fungi | Ascomycota | *Chaetomium* | *NA* | 1 Year |
| 21.29 | 9E-11 | Fungi | Basidiomycota | *Naganishia* | *NA* | 1 Year |
| 21.12 | 1E-10 | Fungi | Ascomycota | *NA* | *NA* | 1 Year |
| 9.51 | 6E-04 | Fungi | Ascomycota | *Collariella* | *Collariella anguipilia* | 1 Year |
| 8.81 | 4E-03 | Fungi | Ascomycota | *NA* | *NA* | 1 Year |
| 8.55 | 1E-04 | Fungi | Ascomycota | *Fusarium* | *NA* | 1 Year |
| 8.23 | 5E-03 | Fungi | Ascomycota | *Clarireedia* | *Clarireedia bennettii* | 1 Year |
| 7.58 | 7E-03 | Fungi | Ascomycota | *Preussia* | *Preussia polymorpha* | 1 Year |
| 6.76 | 3E-04 | Fungi | Ascomycota | *Oedocephalum* | *NA* | 1 Year |
| 5.43 | 3E-04 | Fungi | Ascomycota | *NA* | *NA* | 1 Year |
| 4.93 | 2E-04 | Fungi | Basidiomycota | *Naganishia* | *Naganishia albida* | 1 Year |
| 26.61 | 4E-16 | Fungi | Ascomycota | *NA* | *NA* | 3 Years |
| 25.70 | 4E-15 | Fungi | Ascomycota | *Laburnicola* | *Laburnicola sp* | 3 Years |
| 25.65 | 3E-15 | Fungi | Ascomycota | *Iodophanus* | *Iodophanus sp* | 3 Years |
| 24.46 | 5E-19 | Fungi | Ascomycota | *Cladorrhinum* | *Cladorrhinum sp* | 3 Years |
| 24.29 | 3E-19 | Fungi | Ascomycota | *NA* | *NA* | 3 Years |
| 24.27 | 1E-13 | Fungi | Ascomycota | *Spooneromyces* | *Spooneromyces laeticolor* | 3 Years |
| 24.24 | 1E-13 | Fungi | Ascomycota | *NA* | *NA* | 3 Years |
| 24.15 | 1E-13 | Fungi | Ascomycota | *NA* | *NA* | 3 Years |
| 23.94 | 2E-13 | Fungi | Ascomycota | *Macrophomina* | *NA* | 3 Years |
| 23.77 | 3E-13 | Fungi | Ascomycota | Capnodiales inc. sed. | Capnodiales sp | 3 Years |
| 23.73 | 3E-13 | Fungi | Basidiomycota | *Geastrum* | *Geastrum sp* | 3 Years |
| 23.69 | 3E-13 | Fungi | Ascomycota | *Chaetosphaeronema* | *Chaetosphaeronema sp* | 3 Years |
| 23.66 | 2E-18 | Fungi | Ascomycota | *Edenia* | *Edenia gomezpompae* | 3 Years |
| 23.65 | 3E-16 | Fungi | Ascomycota | *NA* | *NA* | 3 Years |
| 23.59 | 4E-13 | Fungi | Basidiomycota | *Naganishia* | *NA* | 3 Years |
| 23.55 | 4E-13 | Fungi | Ascomycota | *NA* | *NA* | 3 Years |
| 23.32 | 7E-13 | Fungi | Basidiomycota | *Coprinus* | *Coprinus phaeopunctatus* | 3 Years |
| 23.30 | 7E-13 | Fungi | Ascomycota | *Preussia* | *NA* | 3 Years |
| 22.92 | 3E-13 | Fungi | NA | *NA* | *NA* | 3 Years |
| 22.82 | 1E-20 | Fungi | Ascomycota | *Pseudogymnoascus* | *Pseudogymnoascus pannorum* | 3 Years |
| 22.79 | 4E-17 | Fungi | Ascomycota | *NA* | *NA* | 3 Years |
| 22.52 | 3E-15 | Fungi | Ascomycota | *Sulcatistroma* | *Sulcatistroma nolinae* | 3 Years |
| 22.51 | 4E-12 | Fungi | Chytridiomycota | Lobulomycetales inc. sed. | Lobulomycetales sp | 3 Years |
| 22.44 | 1E-16 | Fungi | Ascomycota | *Paraphoma* | *Paraphoma sp* | 3 Years |
| 22.44 | 5E-12 | Fungi | Ascomycota | *Knufia* | *NA* | 3 Years |
| 22.23 | 4E-12 | Fungi | Ascomycota | *NA* | *NA* | 3 Years |
| 22.09 | 1E-19 | Fungi | Ascomycota | *NA* | *NA* | 3 Years |
| 21.97 | 3E-16 | Fungi | Ascomycota | *Gibberella* | *Gibberella avenacea* | 3 Years |
| 21.97 | 6E-13 | Fungi | Ascomycota | *Aspergillus* | *Aspergillus ruber* | 3 Years |
| 21.97 | 1E-11 | Fungi | Basidiomycota | *Naganishia* | *Naganishia albida* | 3 Years |
| 21.95 | 1E-11 | Fungi | Ascomycota | *Gymnoascus* | *Gymnoascus reessii* | 3 Years |
| 21.94 | 1E-11 | Fungi | Ascomycota | *Knufia* | *NA* | 3 Years |
| 21.92 | 1E-11 | Fungi | Basidiomycota | *Panaeolus* | *Panaeolus fimicola* | 3 Years |
| 21.89 | 2E-11 | Fungi | Ascomycota | *Darksidea* | *Darksidea alpha* | 3 Years |
| 21.88 | 2E-11 | Fungi | Ascomycota | *Penicillium* | *NA* | 3 Years |
| 21.87 | 7E-13 | Fungi | Ascomycota | *Westerdykella* | *Westerdykella sp* | 3 Years |
| 21.86 | 1E-14 | Fungi | Ascomycota | *Dactylonectria* | *Dactylonectria macrodidyma* | 3 Years |
| 21.82 | 2E-11 | Fungi | Ascomycota | *Darksidea* | *Darksidea alpha* | 3 Years |
| 21.73 | 2E-11 | Fungi | Ascomycota | *NA* | *NA* | 3 Years |
| 21.67 | 2E-11 | Fungi | Ascomycota | *Coniosporium* | *Coniosporium sp* | 3 Years |
| 21.66 | 2E-11 | Fungi | Ascomycota | *Microdochium* | *Microdochium sp* | 3 Years |
| 21.62 | 2E-11 | Fungi | Ascomycota | *Coniochaeta* | *NA* | 3 Years |
| 21.52 | 8E-13 | Fungi | Ascomycota | *Coniothyrium* | *Coniothyrium sp* | 3 Years |
| 21.41 | 4E-11 | Fungi | Ascomycota | *Acrophialophora* | *NA* | 3 Years |
| 21.41 | 1E-13 | Fungi | Ascomycota | *Podospora* | *Podospora minicauda* | 3 Years |
| 21.39 | 5E-12 | Fungi | Ascomycota | *Schizothecium* | *NA* | 3 Years |
| 20.91 | 8E-12 | Fungi | Ascomycota | *Ascobolus* | *Ascobolus sp* | 3 Years |
| 20.60 | 2E-10 | Fungi | Ascomycota | *NA* | *NA* | 3 Years |
| 20.28 | 2E-11 | Fungi | Ascomycota | *Scutellinia* | *Scutellinia sp* | 3 Years |
| 11.22 | 2E-04 | Fungi | Ascomycota | *Collariella* | *Collariella anguipilia* | 3 Years |
| 9.15 | 2E-03 | Fungi | Ascomycota | *NA* | *NA* | 3 Years |
| 8.99 | 4E-04 | Fungi | Basidiomycota | *Geastrum* | *Geastrum sp* | 3 Years |
| 8.21 | 2E-03 | Fungi | Mucoromycota | *Rhizopus* | *Rhizopus arrhizus* | 3 Years |
| 8.06 | 3E-03 | Fungi | Ascomycota | *Preussia* | *Preussia terricola* | 3 Years |
| 7.22 | 2E-05 | Fungi | Ascomycota | *Iodophanus* | *Iodophanus sp* | 3 Years |
| 6.84 | 1E-02 | Fungi | Basidiomycota | *Naganishia* | *Naganishia randhawae* | 3 Years |
| 6.20 | 1E-04 | Fungi | Ascomycota | *Aureobasidium* | *NA* | 3 Years |
| 4.94 | 2E-08 | Fungi | Basidiomycota | *Naganishia* | *Naganishia albida* | 3 Years |
| 4.85 | 3E-07 | Fungi | Ascomycota | *Sulcatistroma* | *Sulcatistroma nolinae* | 3 Years |
| 4.28 | 1E-11 | Fungi | Ascomycota | *Alternaria* | *NA* | 3 Years |
| 3.43 | 2E-03 | Fungi | Basidiomycota | *Naganishia* | *NA* | 3 Years |
| 1.46 | 2E-03 | Fungi | Ascomycota | *Alternaria* | *NA* | 3 Years |

Table S14. Fungal genera showing negative differential abundance in burned plots (decrease). All rows showed significant decreases in burned plots based on the adjusted p value (padj < 0.01) after a Benjamini–Hochberg correction in DESEQ2.

| log2Fold Change | padj | Kingdom | Phylum | Genus | Species | Timepoint |
| --- | --- | --- | --- | --- | --- | --- |
| -7.58 | 1E-03 | Fungi | Ascomycota | Pezizaceae inc. sed. | Pezizaceae sp | 17 Days |
| -9.20 | 2E-06 | Fungi | Ascomycota | *NA* | *NA* | 17 Days |
| -9.78 | 5E-05 | Fungi | Ascomycota | *Didymella* | *NA* | 17 Days |
| -23.27 | 1E-12 | Fungi | Ascomycota | *Coniozyma* | *Coniozyma leucospermi* | 17 Days |
| -23.29 | 1E-12 | Fungi | Ascomycota | Capnodiales inc. sed. | Capnodiales sp | 17 Days |
| -23.80 | 4E-13 | Fungi | Basidiomycota | *NA* | *NA* | 17 Days |
| -24.82 | 4E-14 | Fungi | Ascomycota | Pezizaceae inc. sed. | Pezizaceae sp | 17 Days |
| -25.14 | 5E-19 | Fungi | Ascomycota | *Knufia* | *NA* | 17 Days |
| -25.67 | 3E-08 | Fungi | Ascomycota | Arthoniomycetes inc. sed. | Arthoniomycetes sp | 1 Month |
| -25.88 | 2E-08 | Fungi | Ascomycota | Pezizaceae inc. sed. | Pezizaceae sp | 1 Month |
| -6.57 | 3E-03 | Fungi | Ascomycota | Pezizaceae inc. sed. | Pezizaceae sp | 8 Months |
| -7.32 | 9E-03 | Fungi | Ascomycota | *Oedocephalum* | *Oedocephalum sp* | 8 Months |
| -7.51 | 6E-03 | Fungi | Ascomycota | Arthoniomycetes inc. sed. | Arthoniomycetes sp | 8 Months |
| -23.93 | 5E-13 | Fungi | Ascomycota | *Phaeococcomyces* | *Phaeococcomyces sp* | 8 Months |
| -24.12 | 3E-13 | Fungi | Ascomycota | *NA* | *NA* | 8 Months |
| -24.26 | 3E-13 | Fungi | Ascomycota | *Libertasomyces* | *Libertasomyces myopori* | 8 Months |
| -24.89 | 6E-14 | Fungi | Ascomycota | *Aureobasidium* | *Aureobasidium iranianum* | 8 Months |
| -25.80 | 2E-23 | Fungi | Ascomycota | Arthoniomycetes inc. sed. | Arthoniomycetes sp | 8 Months |
| -8.14 | 4E-03 | Fungi | Ascomycota | *Ramimonilia* | *Ramimonilia apicalis* | 1 Year |
| -21.88 | 2E-11 | Fungi | Basidiomycota | *NA* | *NA* | 1 Year |
| -24.18 | 1E-13 | Fungi | Ascomycota | *Ascobolus* | *Ascobolus sp* | 1 Year |
| -24.19 | 1E-13 | Fungi | Ascomycota | *NA* | *NA* | 1 Year |
| -24.46 | 1E-13 | Fungi | Ascomycota | Pezizaceae inc. sed. | Pezizaceae sp | 1 Year |
| -24.53 | 1E-13 | Fungi | Ascomycota | *Ascobolus* | *Ascobolus sp* | 1 Year |
| -24.81 | 6E-14 | Fungi | Ascomycota | *Kalmusia* | *Kalmusia utahensis* | 1 Year |
| -25.51 | 1E-14 | Fungi | Ascomycota | *NA* | *NA* | 1 Year |
| -8.35 | 1E-05 | Fungi | Ascomycota | *Dematiopleospora* | *Dematiopleospora sp* | 3 Years |
| -8.69 | 2E-04 | Fungi | Ascomycota | *Pyrenophora* | *Pyrenophora sieglingiae* | 3 Years |
| -23.33 | 5E-13 | Fungi | Ascomycota | *NA* | *NA* | 3 Years |
| -23.50 | 3E-13 | Fungi | Ascomycota | *Helicoma* | *Helicoma sp* | 3 Years |
| -23.85 | 2E-13 | Fungi | Ascomycota | Pezizaceae inc. sed. | Pezizaceae sp | 3 Years |
| -25.03 | 1E-14 | Fungi | Ascomycota | *Helicoma* | *Helicoma sp* | 3 Years |
| -25.41 | 4E-15 | Fungi | Ascomycota | *NA* | *NA* | 3 Years |
| -25.59 | 9E-19 | Fungi | Ascomycota | Capnodiales inc. sed. | Capnodiales sp | 3 Years |
| -25.94 | 1E-15 | Fungi | Ascomycota | Chaetomiaceae inc. sed. | Chaetomiaceae sp | 3 Years |

Figure S1. Plant species richness at each of the 9 plots (6 burned plots in red and 3 unburned plots in blue) in September 2020, 2021, and 2023 corresponding to 17 days, 1 year and 3 years post-fire.

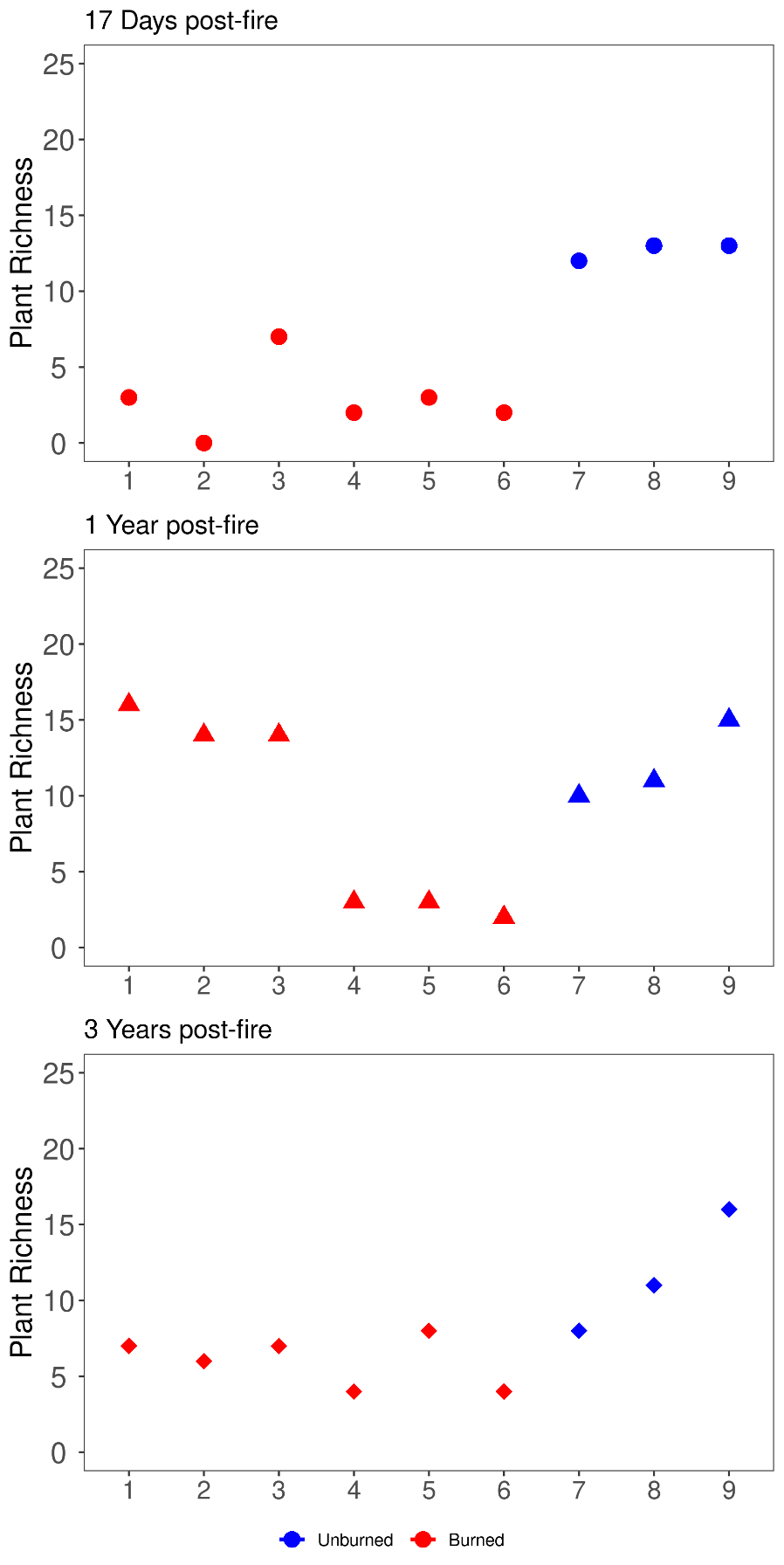

Figure S2. Soil characteristics at available timepoints indicating per plot means (center point) and standard error (bars) as well as data points (small points) for A) pH with a dashed line indicating pH 7, B) phosphate, C) ammonium, and D) nitrate and nitrite concentrations in unburned (blue) versus burned (red) plots. Asterisks indicate significant differences between burned and unburned means at each time point based on generalized mixed effects models for each variable.

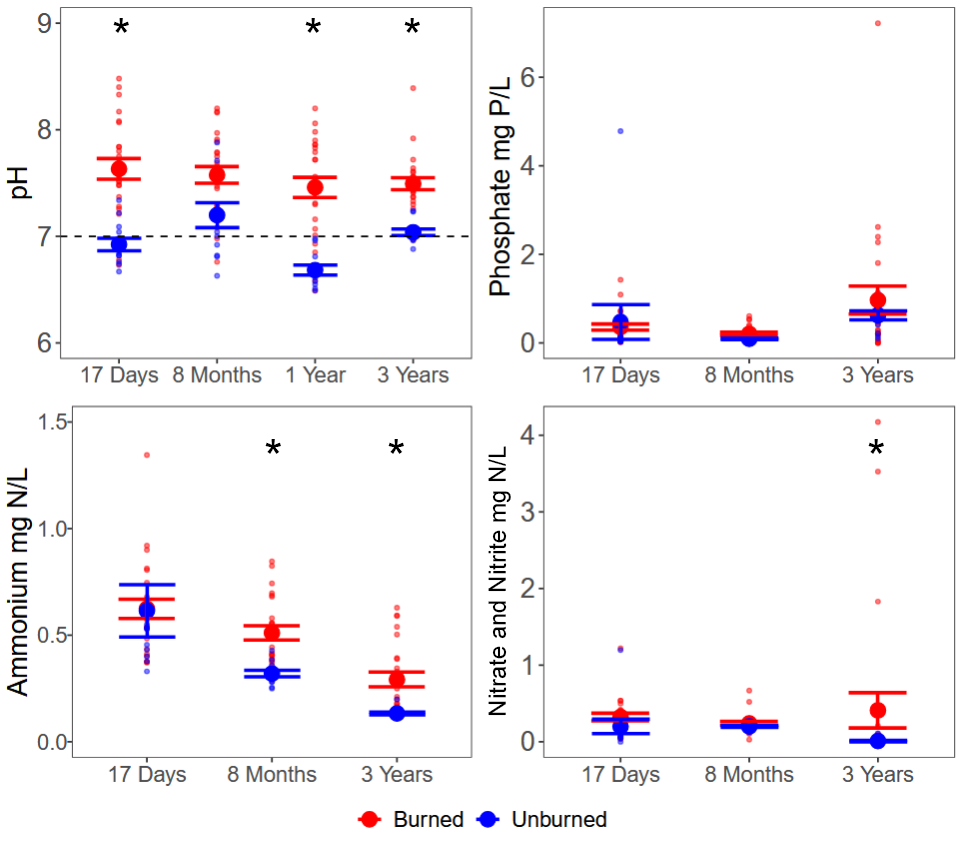

Figure S3. Fungal richness estimated by number of fungal amplicon sequence variants (AVS) regressed against ash depth as measured at 17-days post-fire. We tested richness and ash depth as a linear model with R^2^ and p-values listed per comparison.

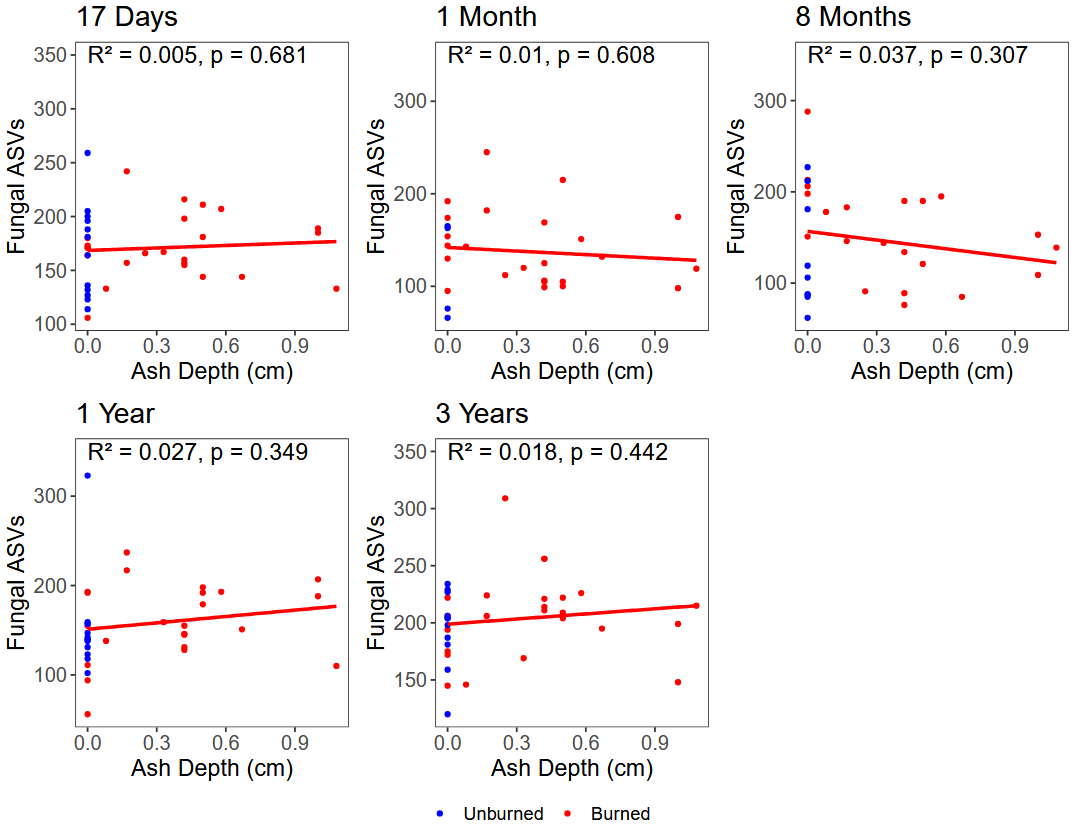

Figure S4. Bacterial richness estimated by number of bacterial amplicon sequence variants (AVS) regressed against ash depth as measured at 17-days post-fire. We tested richness and ash depth as a linear model with R^2^ and p-values listed per comparison.

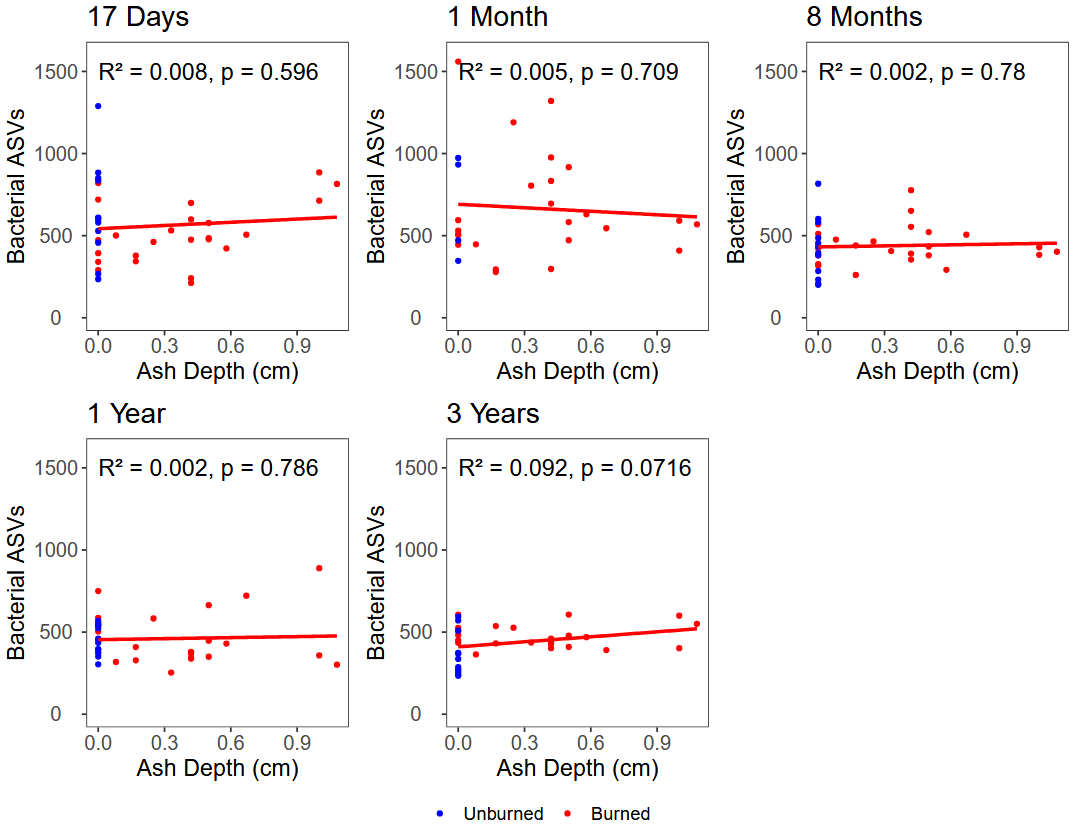

Figure S5. Relative sequence abundance of archaeal and bacterial A) phyla and B) genera with each timepoint colored in blue for all unburned plots and red for all burned plots.

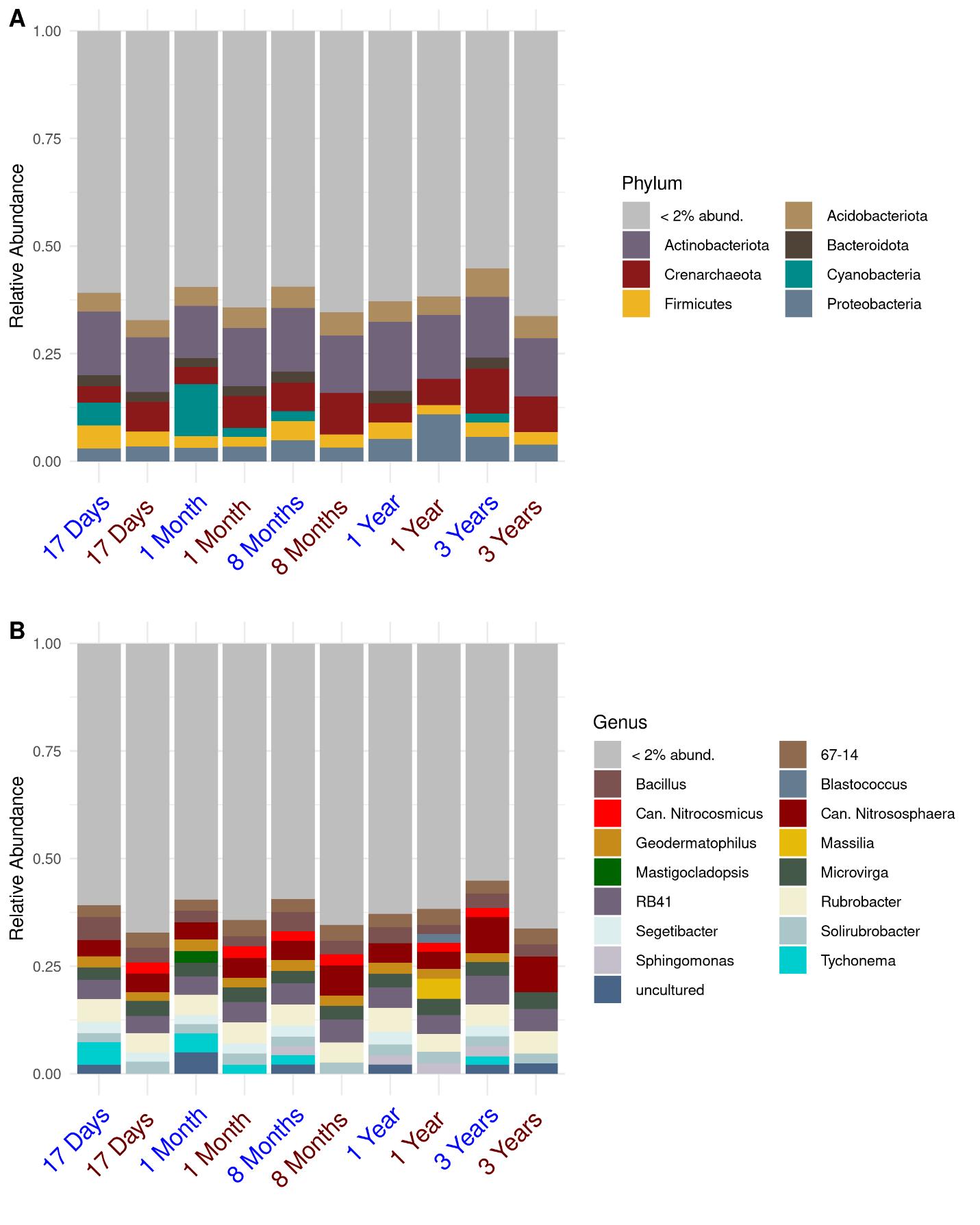

Figure S6. Relative sequence abundance of fungal A) phyla and B) genera at each timepoint colored in blue for all unburned plots and red for all burned plots.

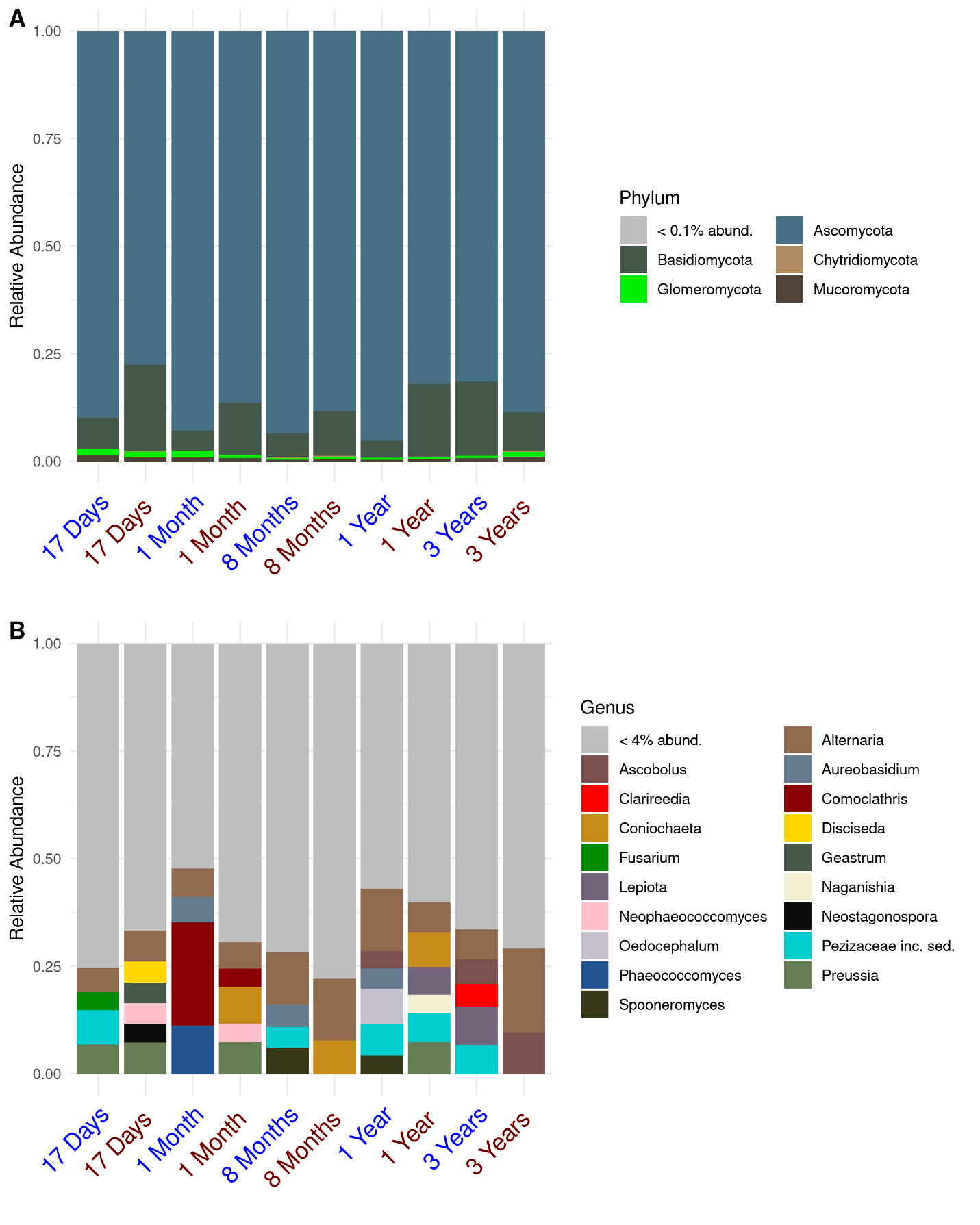

Figure S7. Glomeromycotina relative sequence abundance in burned and unburned plots at each timepoint.

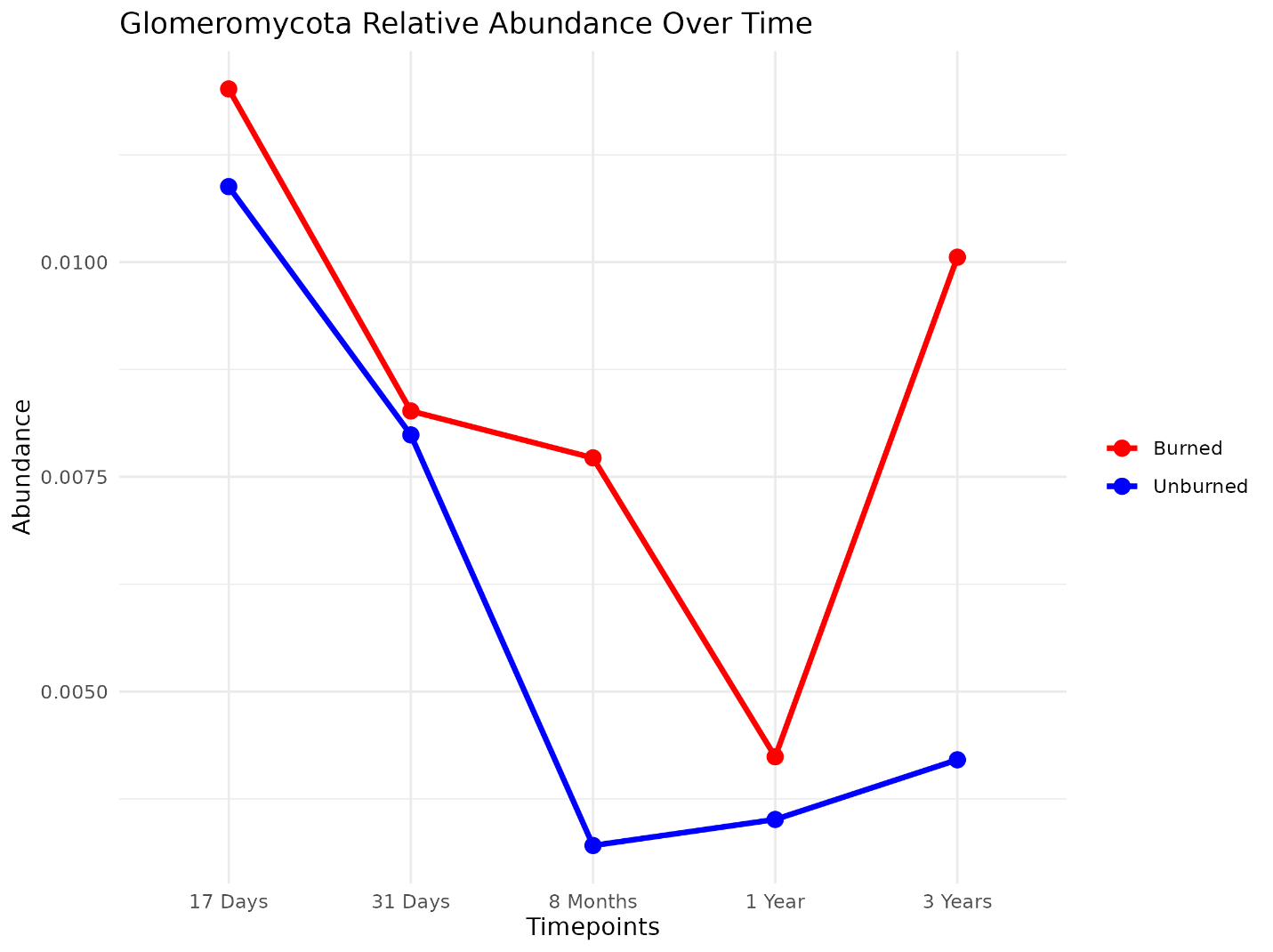

Figure S8. Fungal guild relative sequence abundance over time. Guilds represented primarily by one genus have the genus listed below the guild name. Darker lines indicate relative abundance in burned plots. Light lines indicate relative abundance in unburned plots.

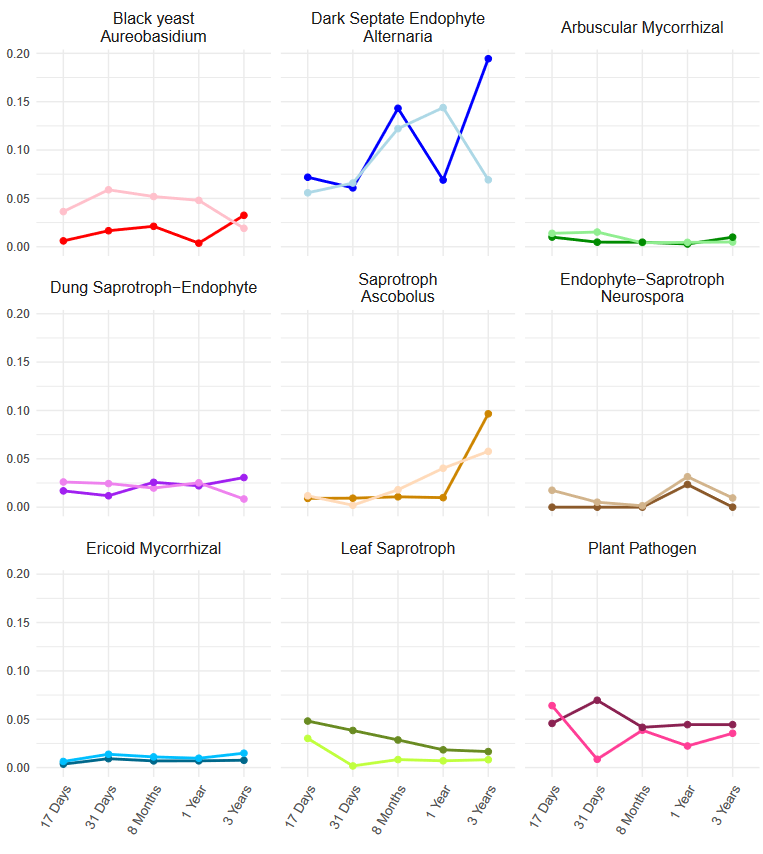
